# CHARR efficiently estimates contamination from DNA sequencing data

**DOI:** 10.1101/2023.06.28.545801

**Authors:** Wenhan Lu, Laura D. Gauthier, Timothy Poterba, Edoardo Giacopuzzi, Julia K. Goodrich, Christine R. Stevens, Daniel King, Mark J. Daly, Benjamin M. Neale, Konrad J. Karczewski

## Abstract

DNA sample contamination is a major issue in clinical and research applications of whole genome and exome sequencing. Even modest levels of contamination can substantially affect the overall quality of variant calls and lead to widespread genotyping errors. Currently, popular tools for estimating the contamination level use short-read data (BAM/CRAM files), which are expensive to store and manipulate and often not retained or shared widely. We propose a new metric to estimate DNA sample contamination from variant-level whole genome and exome sequence data, CHARR, Contamination from Homozygous Alternate Reference Reads, which leverages the infiltration of reference reads within homozygous alternate variant calls. CHARR uses a small proportion of variant-level genotype information and thus can be computed from single-sample gVCFs or callsets in VCF or BCF formats, as well as efficiently stored variant calls in Hail VDS format. Our results demonstrate that CHARR accurately recapitulates results from existing tools with substantially reduced costs, improving the accuracy and efficiency of downstream analyses of ultra-large whole genome and exome sequencing datasets.

## Introduction

Cohorts of individuals with WGS and WES data continue to grow exponentially, necessitating scalable and efficient methods to perform quality control and analysis. One of the most potentially pernicious issues during the generation of sequencing data is DNA sample contamination, in which the DNA of the target sample is mixed with DNA from external samples. Even slight contamination can affect sequencing accuracy and introduce biases in downstream variant analysis and association tests.

There are several existing methods to detect DNA sample contamination. ContEst (Cibulskis et al., 2011) maximizes a posteriori estimate of contamination based on the base identities and quality scores from VCF and BAM files. VerifyBamID, a popular tool to identify sample contamination, estimates the contamination level by maximizing the likelihood constructed from a two-sample mixture model with sequencing data and external genotype information (Jun et al., 2012). A recent update, VerifyBamID2, jointly estimates the contamination rate and genetic ancestry in a unified likelihood framework without assuming known population allele frequencies (Zhang et al., 2020). Conpair, focusing on contamination in tumor samples, uses BAM files, the reference genome, and a short list of pre-selected highly informative genomic markers to estimate contamination from a probabilistic model generalized from VerifyBamID (Bergmann et al., 2016). Peddy uses a VCF file with an associated PED/FAM file to infer contamination level by investigating the properties of heterozygous calls (Pedersen & Quinlan, 2017). These tools have successfully detected contamination levels in many sequencing data studies with high accuracy and sensitivity. However, they either operate on the underlying short-read data (CRAMs or BAMs), which is computationally expensive, or do not directly estimate contamination rates. With sample sizes soon reaching millions of individuals with WGS, even pipelines that cost tens of cents per sample will incur high aggregate costs. As increasing large-scale WGS and WES data are processed into the gVCF format, a gVCF-based metric for contamination estimation would avoid the heavy computational burden of CRAM/BAM-based analysis.

We have previously discovered a systematic undercalling of homozygous genotypes, particularly at common indel sites and identified the cause as a genotyping error mode whereby contaminating reference alleles result in erroneous heterozygous calls with high allele balance (Karczewski et al., 2019). Based on this empirical observation and recent optimization of genotype data structures, we propose a novel metric, CHARR, Contamination from Homozygous Alternate Reference Reads, to estimate contamination level from genotype data without requiring raw read data. As the name suggests, it evaluates contamination by investigating the level of observed reference reads within homozygous alternate variant calls. CHARR only requires a very basic set of genotype information, and it does not rely on data from reference blocks. It thus can be computed from single-sample gVCFs or cohorts in VCF or BCF formats. With the newly designed Hail format for storing variant call data, VariantDataset (VDS), which enables more flexible operations on the genotype data by splitting variant calls and reference blocks into two component MatrixTables, CHARR can be conveniently run using only the variant data component of the VDS. With substantially reduced time and cost in computation, CHARR can be used to conveniently identify samples with low sequencing quality at early stages of sequencing studies, which will facilitate the implementation of stringent quality control of samples and improve the accuracy and efficiency of variant discovery in ultra-large sequencing datasets in the future.

### Pervasive infiltration of reference reads among variants in gnomAD

With high-quality data, heterozygous and homozygous variants should, in expectation, have 50% and 0% allelic contribution from reference alleles. However, the presence of contamination can affect these estimates, and indeed in some homozygous variants, we observe the presence of reference reads (Figure 1). We performed a systematic analysis of homozygous variants in gnomAD v3 (defined as those autosomal bi-allelic SNVs with GQ ≥20 and 20 ≤ DP ≤ 100 -mean of 584K per sample) and found that all 59,765 assayed WGS samples have at least one high-quality homozygous variant with a reference allele balance over 10% and 51,849 have more than 10,000 high-quality homozygous variants with reference reads observed (Figure S1 & Table S1). Using these genotypes, we derived a new contamination estimator, CHARR, which evaluates the sample-wise mean proportion of reference reads of homozygous alternate variants adjusted by their corresponding allele frequencies (Methods).

**Figure 1.**
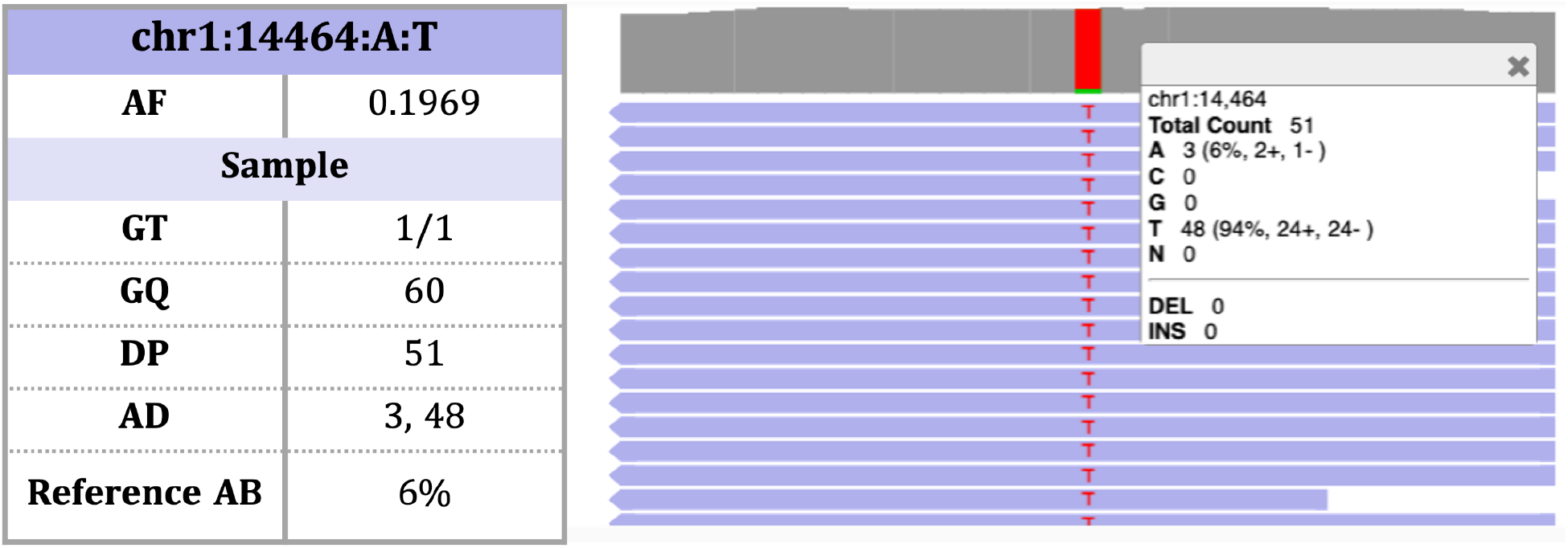
An example of a homozygous variant in gnomAD v3 with infiltration of reference reads. For this specific sample, the variant has a genotype quality (GQ) of 60 and a depth (DP) of 51, indicating a high-quality genotype. However, we observe a relatively high proportion of reference reads (AD[0]/DP = 3/51 = 6%), suggesting the presence of potential contamination.

### Concordance with Freemix Score in gnomAD

We evaluated CHARR on 59,765 whole genome samples from gnomAD v3 including 58,986 samples sequenced at the Broad Institute and 779 from the HGDP callset (Figure 2A). Both platforms demonstrated strong correlations (r_Broad_ = 0.988, p_Broad_ < 10^−100^; r_HGDP_ = 0.967, p_HGDP_ < 10^−100^) between CHARR and the Freemix score from VerifyBamID (Figure S3A). In 102,063 WES samples from gnomAD v2, despite having fewer homozygous variants (Figure 2B), CHARR was still highly correlated (r_Broad_=0.990, p_Broad_ < 10^−100^; r_Other_=0.987, p_Other_ < 10^−100^) with the Freemix score (Figure S3B). We repeated this analysis with several different filtering parameters to ensure the robustness of the results. In particular, we investigated cutoffs for ref-AF (0%, 1%, 10%, and 20%), and for sequencing depth (None, 10x-100x, and 20x-100x), as well as selections of variant types (all, SNPs, and Indels). We find that our method is robust to these choices in parameters, but suggest a series of cutoffs for optimal analysis (see Methods). We also find that this method is not biased by somatic mosaicism as we observe no correlation with age and robustness to the removal of known mosaic genes (Figures S7-9, Table S2).

**Figure 2.**
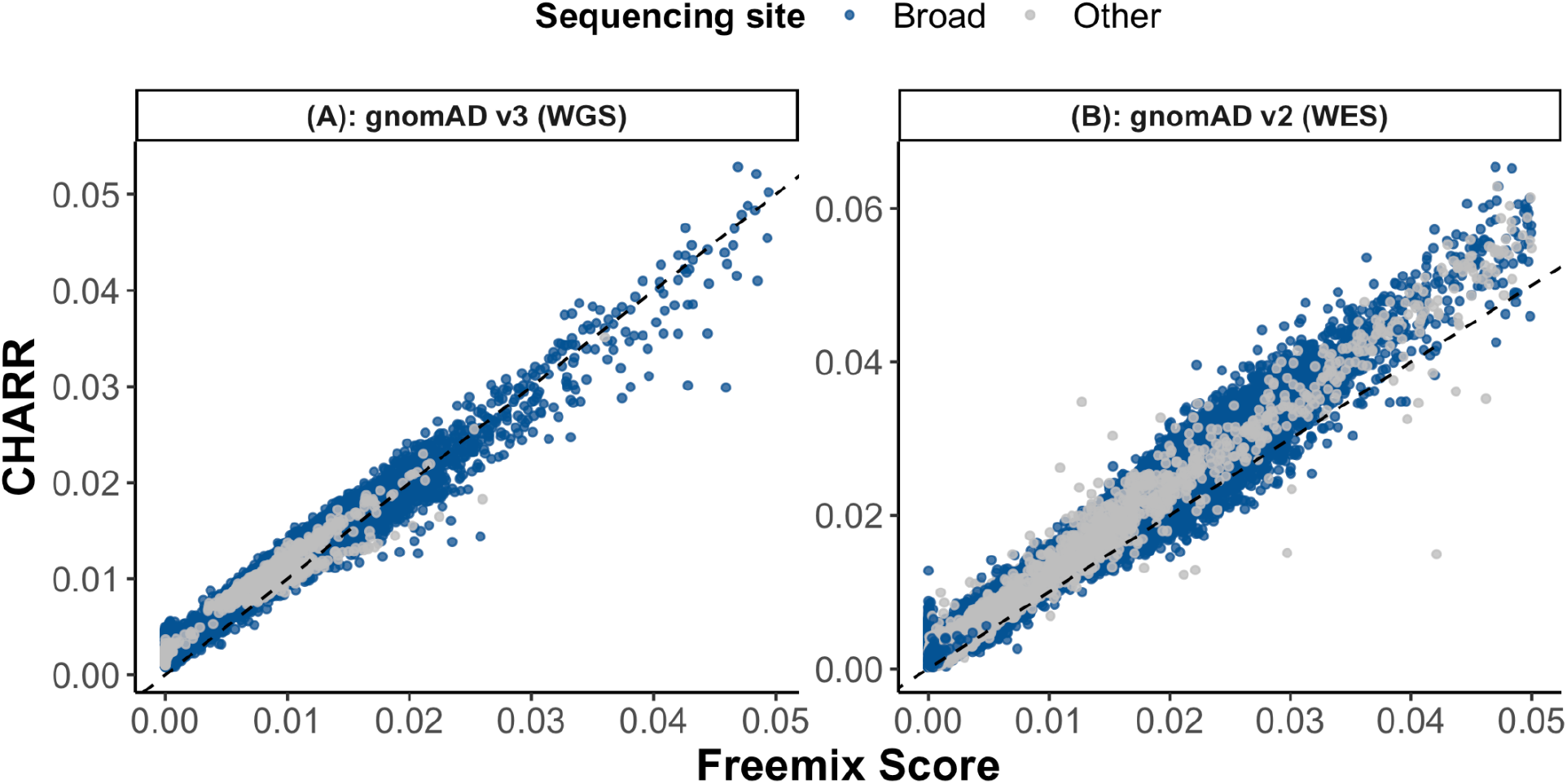
A comparison of Freemix score and CHARR for 59,765 gnomAD v3 release WGS and 102,063 v2 release WES samples. The mean numbers of homozygous variants used for computing CHARR are 463668 and 718, respectively for (A) and (B). The black dashed line represents y=x.

### Simulation results demonstrate mechanisms of contamination

To produce a ground truth for comparison, we implemented a simulation pipeline to create samples with known contamination levels. First, we randomly selected 30 samples from the HGDP dataset distributed across 6 genetic ancestry groups and the spectrum of contamination level (Figure S12). Based on this subset, we built a decontamination pipeline to remove the existing contamination in each of the 30 files, which replaces the reads in the CRAM files of the samples with the corresponding alleles in the reference genome and inserts the ‘true’ variants observed from the gVCF files of the samples (Methods: CRAM file decontamination; Figure S13). From these 30 contamination-free CRAM files, we applied our n-way mixing simulation pipeline at five selected contamination rates: 0.5%, 1%, 2%, 5%, and 10%, and compared the performance of CHARR and VerifyBamID on 150 simulated CRAM files with diverse sources of contaminant (Methods: Mixing N contamination-free samples).

CHARR and VerifyBamID produce consistent results across the five contamination levels under the n-way mixing simulation (r = 0.997, p = 3.45×10^−169^). However, CHARR evaluates contamination levels more accurately than VerifyBamID across all true contamination rates (Figure 3). At high contamination levels (10%), we observe a slight deflation in both methods: on average, CHARR is deflated by 10%, while VerifyBamID is deflated by 25%.

**Figure 3.**
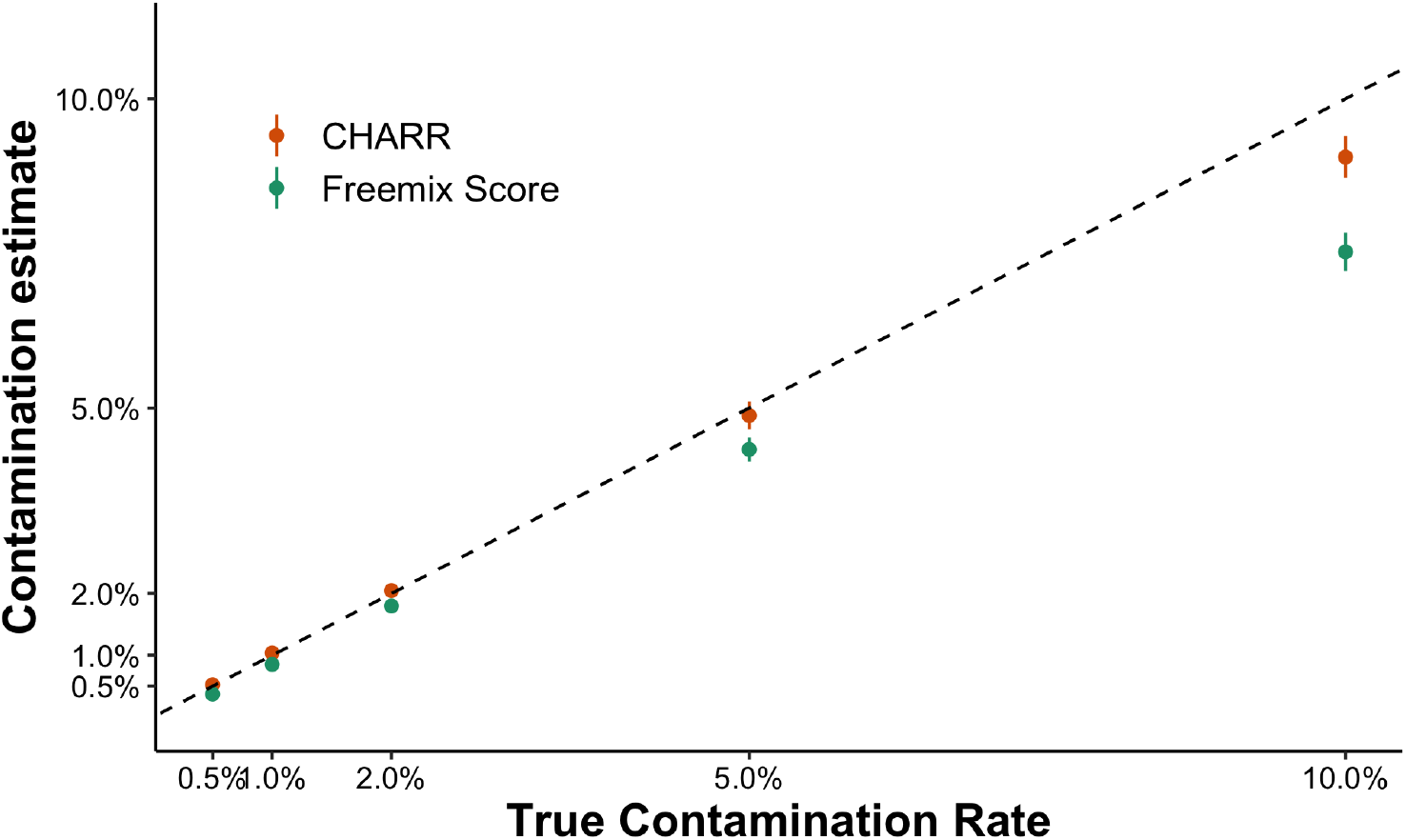
A comparison of Freemix score (green) and CHARR (orange) for 150 simulated n-way mixed samples across five true contamination levels (x-axis). CHARR is computed using the local allele frequencies filtered to variants with 100% callrate among the 30 decontaminated samples. The black dashed line represents y=x.

### Computational efficiency

A major advantage of CHARR stems from its efficiency by using only non-reference variant call data, which can be obtained from VCF, gVCF, or our latest VariantDataSet (VDS) structure, without requiring individual-level BAM or CRAM files. We have implemented CHARR on data from hundreds of thousands of samples using the optimized, scalable VDS format. By using only the variant call component MatrixTable of the VDS, it takes 2,150 CPU-hours to compute CHARR for all 153K WGS samples in gnomAD v3 (regardless of release status), amortizing to a cost of $280 per 1M samples. It also accurately recapitulated the contamination levels for the 103k WES samples from gnomAD v2 in 250 CPU-hours, for a cost of $48 per 1M samples. The computation of CHARR is more than two orders of magnitude cheaper than running the VerifyBamID on CRAM files, which takes about 45 mins per sample (672 CPU-hours in total) and costs $155 to compute Freemix scores for the 948 WGS samples from HGDP (Table 1). Based on this comparison and the previous results, it is feasible to downsample whole genome data to the exonic regions to control for computational cost while still achieving a reasonable estimate of DNA sample contamination with CHARR.

**Table 1.** Comparison of the cost and computational efficiency of VerifyBamID and CHARR

|  | VerifyBamID | CHARR |  |
| --- | --- | --- | --- |
| Source | HGDP | gnomAD v3 | gnomAD v2 |
| Data | WGS | WGS | WES |
| Input format | CRAM | VDS | VDS |
| N samples | 948 | 153,030 | 103,017 |
| Total runtime (CPU-hours) | 672 | 2,150 | 250 |
| Runtime per sample (CPU-hours) | 0.709 | 0.014 | 0.002 |
| Total costs (USD) | \$155 | \$43 | \$5 |
| Cost per 1M samples (USD) | \$160,000 | \$280 | \$48 |

## Discussion

Here, we have described a novel metric to evaluate the contamination level of DNA samples from variant calls with high accuracy and efficiency. There are a few considerations and limitations to our work. In particular, using the reference allele frequency that best reflects the structure of the contaminating samples, which is inherently unknown, is important for accurately determining CHARR. The best practice may be to obtain the allele frequency information from the callset directly; however, this is often not ideal due to insufficient sample size or may not accurately reflect the contamination sources. We have applied two different sources of allele frequency to compute CHARR on the gnomAD v2 WES set (Figures S5 & S6), where results from both sources present high correlations (r > 0.98, p < 10^−100^) with the corresponding Freemix scores. Therefore, we recommend direct use of allele frequencies from the gnomAD data, especially when the local sample size is small, which approximates contamination rates well, avoids additional computation, and has no access restrictions (Supplement: Sources of reference allele frequencies). To address potential influence of genetic mosaicism on our results, we examined the relationship between CHARR and age, which positively correlates with the occurrence of mosaicism, and found no evidence of correlation across all age groups within both the WGS and the WES data (Table S2 & Figure S9).

One challenge highlighted by our study is the estimation of high levels of contamination, particularly those that typically fail quality control in large sequencing studies (>5-10%). In these cases, excessive infiltration of reference reads will convert homozygous alternate variant sites to heterozygous, decreasing the variants contributing to CHARR and leading to deflation in contamination estimates. As demonstrated in the n-way simulation results (Figure 3), CHARR presents an approximately 10% downward bias when the true contamination rate is 10%.

Accordingly, we recommend computing the ratio of heterozygous to homozygous variants as a sample quality control step to identify potentially highly contaminated samples. However, since a typical quality control protocol will set a contamination cutoff around 5%, CHARR is still capable of capturing the highly contaminated samples that would be filtered out.

Our results yield insights into the dynamics of DNA sample contamination. We find that CHARR is more consistent with VerifyBamID in simulations where multiple samples are the source of contamination rather than a single sample at contamination levels of 5-10% (Figure S17); in the latter case, we observe a deflation likely due to homozygous to heterozygous genotype conversion. This result, coupled with the observation of samples with estimates of ∼5% of both CHARR and VerifyBamID in real data (Figure 2), suggests that contamination from multiple samples is a more likely scenario than single-sample contamination. One limitation of the simulation framework is the relatively small sample size (30 decontaminated samples) compared to typical sample sizes in real-world cohorts (hundreds of thousands of samples). A larger simulation may reveal more precise insights into the mechanisms of contamination.

Overall, CHARR provides an accurate, efficient, and easily accessible solution to estimating DNA sample contamination levels. It can be computed flexibly from any format of variant call data at a wide range of sample size scales, which avoids expensive manipulation on short read data. With extremely low cost and high efficiency, CHARR makes it feasible to estimate contamination levels for ultra-large cohorts, facilitating more scalable quality control processes for sequencing data with millions of samples.

## Methods

### Data availability

We used data from the Genome Aggregation Database, including:

- 59,765 release whole genome samples in gnomAD v3 (Chen et al., 2022)
  - 58,986 samples sequenced at the Broad Institute + 779 HGDP samples
- 102,063 release whole exome samples in gnomAD v2 (Karczewski et al., 2020)
  - 103,027 samples from gnomAD v2, excluding 10 samples with fewer than 10,000 heterozygous variants and 954 samples with an old version of freemix used
- 948 HGDP samples in gnomAD v3 (Koenig et al., 2023)

The WGS samples from gnomAD v3 (Chen et al., 2022) were called individually using the GATK best practices using GATK4 for BQSR and GATK3.5 for HaplotypeCaller to produce gVCFs, and were then jointly called using gVCF combiner in Hail. The WES samples from gnomAD v2 (Karczewski et al., 2020) were called individually using local realignment by GATK HaplotypeCaller (version nightly-2015-07-31-g3c929b0) and were then jointly genotyped for high confidence alleles using GenotypeGVCFs version 3.4-89-ge494930.

### Variant selection

To ensure high and stable qualities of variants, we first filter down to autosomal, bi-allelic SNVs with sequencing depths (DP) of over 20x and below 100x and genotype quality (GQ) of over 20. Since the fraction of reference reads for a homozygous alternate variant is expected to be zero, any significant deviation from these values indicates a potential infiltration of reference reads from external sources, likely resulting from DNA sample contamination. Therefore, we further select the homozygous alternate variants to evaluate contamination levels.

### SVCR: Scalable Variant Call Representation

The most common format for storing large-scale genomic data is the project VCF (multi-sample VCF), a dense format joint called from the gVCF files of all the samples within the set. A downside of this format is that as the sample size increases, the size of the entire data grows exponentially in the order of O(n^1.5^).

To meet the growing demands in processing ultra-large-scale WGS and WES data, we have developed SVCR, the Scalable Variant Call Representation. SVCR is a lossless transformation of gVCFs, which supports incremental/batch merges rather than requiring expensive fresh joint calling pipelines and grows linearly with the increase of sample size. Like the project VCF representation, the scalable variant call representation is a variant-by-sample matrix of records. The fundamental differences between the two representations are 1) a sparse structure that optimizes the storage of reference calls; and 2) a scalable structure for genotype information at multi-allelic sites (Figure M1). With these two features, SVCR reduces the duplication of reference information across samples and avoids the quadratic explosion of genotype information at multi-allelic sites, which provides linear scalability that hugely increases the efficiency of large genomic data presentation. In particular, an implementation of SVCR, the Variant DataSet (VDS), separates the genotype information into two component matrix tables: the reference blocks and the variant calls, which makes it feasible to assess non-reference information without touching large reference blocks, increasing computation efficiency.

As a method to estimate the level of DNA sample contamination, CHARR only requires a very basic set of genotype information. For example, it works with the variant information from a VCF file or any genotype data format that includes sufficient information. As only homozygous genotypes are required, we can run CHARR conveniently with only the variant data part of the VDS, which makes the simple computation process even cheaper.

**Figure M1.**
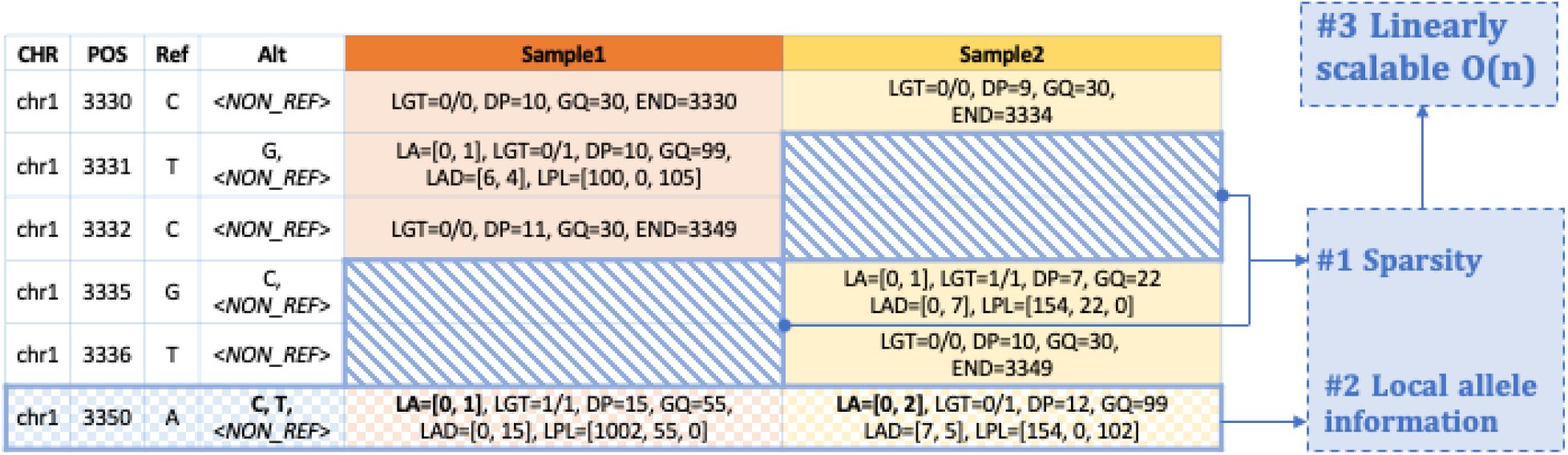
A schematic of the SVCR. Rows indicate variants that exist in at least one sample in the data. The first four columns describe the key information of the variants, and the following ones store the local allele information for each sample. This format scales variant call data linearly by not duplicating allele information for the reference blocks (sparsity; cross-hatched blue) and using locally indexed fields (LA: local alleles, LGT: local genotypes, LAD: local allele depth, LPL: local phred-scaled likelihoods).

### CHARR: a metric to estimate contamination

We define reference allele balance (AB_ref_) as the Allele Depth (AD) of the reference allele divided by the sum of the allele depths of the reference allele and the alternate allele. DNA sample contamination affects the number of reference reads at homozygous alternate sites by:

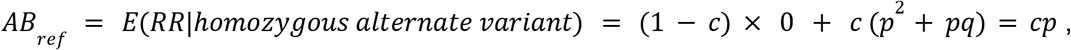

where *c* represents the actual contamination rate and *p, q* are the allele frequencies of the reference allele and the alternate allele respectively. With that, we propose CHARR, Contamination from Homozygous Alternate Reference Reads, as the mean of 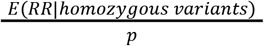 within each sample, 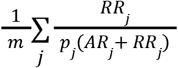 (where *m* is the number of variants included in the calculation), which approximates the level of contamination.

We also constructed the following likelihood function of observing the read data of homozygous variants from *sample* _*i*_. *c*_*i*_ is the contamination rate, and *p* is the reference allele frequency in the population.

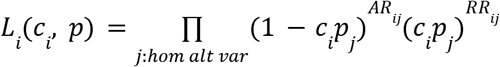

The maximum log-likelihood estimator of the likelihood function, *c* _*mle*_, can be written as 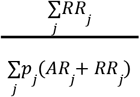 under the condition where 1 − *c*_*i*_ *p*_*j*_ approaches 1, which is generally satisfied. Because *RR*_*j*_ has an approximately linear relationship with *p*_*j*_ (*RR*_*j*_ + *AR*_*j*_), CHARR, written as 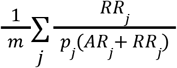, is a close approximation of *c*_*mle*_. To validate CHARR, we performed a grid search on *c* ranging from 0 to 0.1, incremented by 0.001 to maximize the likelihood model. The outcome *c*_*mle*_ is highly aligned with CHARR, with a correlation of 0.996 (p < 10^−100^).

#### Parameter Configuration & Robustness

In the computation of CHARR, we adjust for the abundance of reference alleles among the population by dividing the reference allele balance by its corresponding allele frequency (see CHARR equation above). However, for some homozygous variants, the allele frequency of their corresponding reference alleles is extremely low, which combined with a very low reference allele balance due to sequencing noise will return a very large value of CHARR. As a result, it is not ideal to include homozygous variants with extremely low ref-AF, which will add a disproportionately high value to the overall estimate (Figure S2). Therefore, we have also assessed the score at relative frequency cutoffs of 1%, 5%, 10%, and 20%. For any cutoff at or above 5%, CHARR achieves a high correlation (R-squared > 0.96) with Freemix (Figure S3). Thus, we recommend applying a symmetric cutoff on the reference allele frequency of at least 5% to maintain reasonable accuracy of contamination level.

With a genotype quality filter imposed (GQ ≥ 20), varying cutoffs for sequencing depth does not substantially affect CHARR (Figure S3). We observe similar patterns consistently in both gnomAD v2 WES samples and v3 WGS samples. Moreover, CHARR performs reasonably well with even the biallelic indels, which can be used as an adequate alternate approach with a substantially reduced computational burden (Figure S4). We further show that this estimate is not biased by somatic mosaicism, as it is robust to removal of known mosaic genes and not correlated with age (Figure S7-9, Table S2).

Therefore, in this paper, we refer to CHARR as the contamination estimator computed with autosomal, bi-allelic homozygous alternate SNPs with 20 ≤ DP ≤100, GQ ≥ 20, and 10% < reference AF < 90%, unless otherwise mentioned. However, given the robust performance of CHARR across different combinations of parameter thresholds, all parameter thresholds, including minimum depth, relative frequency, and type of variants, are highly configurable across their spectra. Any sensible selection of these parameters that results in a mean of at least 500 variants is sufficient to detect CHARR under moderate contamination (< 5%) with sufficient accuracy. Further, this result applies to both WGS and WES data, suggesting that it is also feasible to downsample the WGS data to the exonic regions to reduce the computation burden further while achieving an accurate estimate of contamination level.

#### Implementations

We built two implementations of CHARR, one in Hail and one using the VCF format:

- Hail (Hail Team, 2023): hail.methods.qc.compute_charr() (https://github.com/hail-is/hail)
  - The function computes CHARR on Hail MatrixTables and Variant Datasets (VDS), both of which can be generated by importing a VCF file or set of gVCFs
- VCF: https://github.com/HTGenomeAnalysisUnit/SCE-VCF

### Simulation framework

In order to compare the accuracies of CHARR and VerifyBamID, we designed a pipeline to simulate potential contamination scenarios and manually introduce a series of known contamination rates. For each sample, we first decontaminated their short-read data by incorporating information from their corresponding gVCF files and the reference genome. We then simulated contamination using two scenarios: 1) a two-way mixing regime in which we randomly paired the samples and introduced short reads from a contaminating sample to a target sample at a range of contamination rates, and 2) a n-way (in practice, 30-way) regime in which we mixed reads from all decontaminated samples at a range of contamination rates. For each resulting contaminated sample, we then computed contamination estimates using CHARR and VerifyBamID. The entire pipeline is implemented in Hail Batch.

#### CRAM file decontamination

To create a ground truth, we developed a method to remove existing contaminating reads from the short read data before manually introducing a known level of contamination (Figure M2). We randomly selected 30 samples from the HGDP dataset, which included 5 samples from each of the 6 genetic ancestry groups, with their original contamination rates approximately distributed uniformly within each group (Figure S12), and obtained the CRAM files and gVCF files for each sample (Bergström et al., 2020).

From the gVCF files, we identify heterozygous or homozygous alternate SNV calls, excluding indel variants for simplicity. For each read in a given CRAM file, we loop through each position, and check whether a variant is called at this site in the sample’s gVCF file. If so, we randomly sample an allele from the known genotype states (100% for a homozygous alternate allele, 50% for a heterozygous allele) with additional error introduced based on the quality score of that position, and then insert this new allele into the corresponding position. If no variant is present, we insert the corresponding allele in the reference genome. We parallelize this pipeline by chromosome and run GATK HaplotypeCaller on the decontaminated CRAM files to produce decontaminated gVCF files. We then concatenate the chromosome-wise CRAM files into a full CRAM to run VerifyBamID and merge the chromosome-wise gVCFs into full gVCFs. We combine the 30 samples with gVCFs into a VDS and compute CHARR (Figure S13).

From these decontaminated samples, we developed two versions of our contamination simulation pipeline: 1) two-way mixing, and 2) n-way mixing.

**Figure M2.**
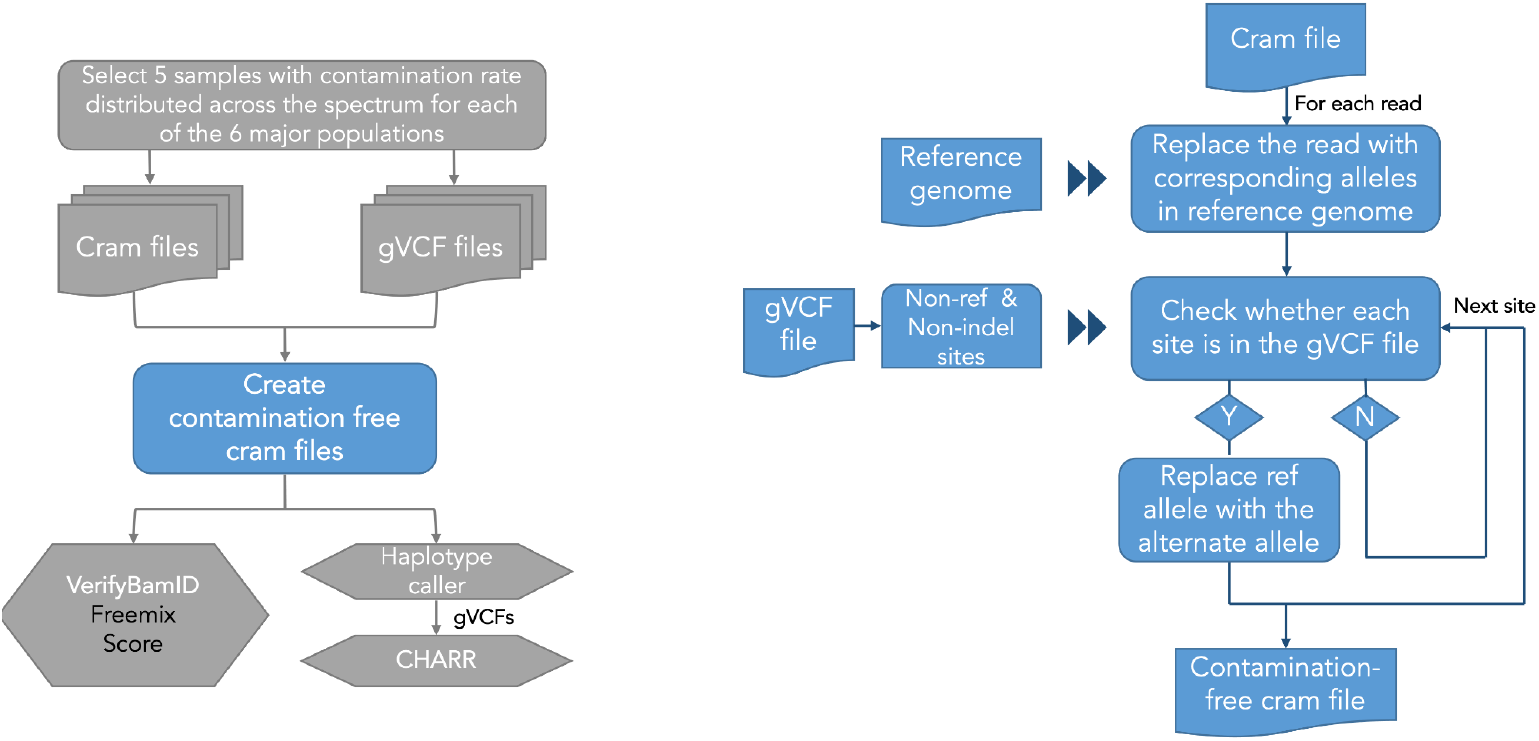
A flowchart describing the pipeline for CRAM file decontamination. Pipeline for computing contamination rate on simulated data (left). Pipeline for decontaminating a single CRAM file (right).

#### Mixing two contamination-free samples

To produce simulated samples with known contamination rates, we started with a simple contaminant and target sample design (Figure M3). Among the 30 samples across 6 populations, we randomly pair a sample (defined as the original sample) with another sample from a different ancestry group (as the contaminant sample) without replacement, and mix the reads from the decontaminated CRAM files at five selected contamination rates: 0.5%, 1%, 2%, 5%, and 10%, thus generating 150 mixed CRAM files each with only a single source of contaminating reads. Within each pair, we pick the first sample as the original sample and the one paired to that sample as the contaminant, so that each sample serves as the original and the contaminant sample both exactly once (Figure M4).

**Figure M3.**
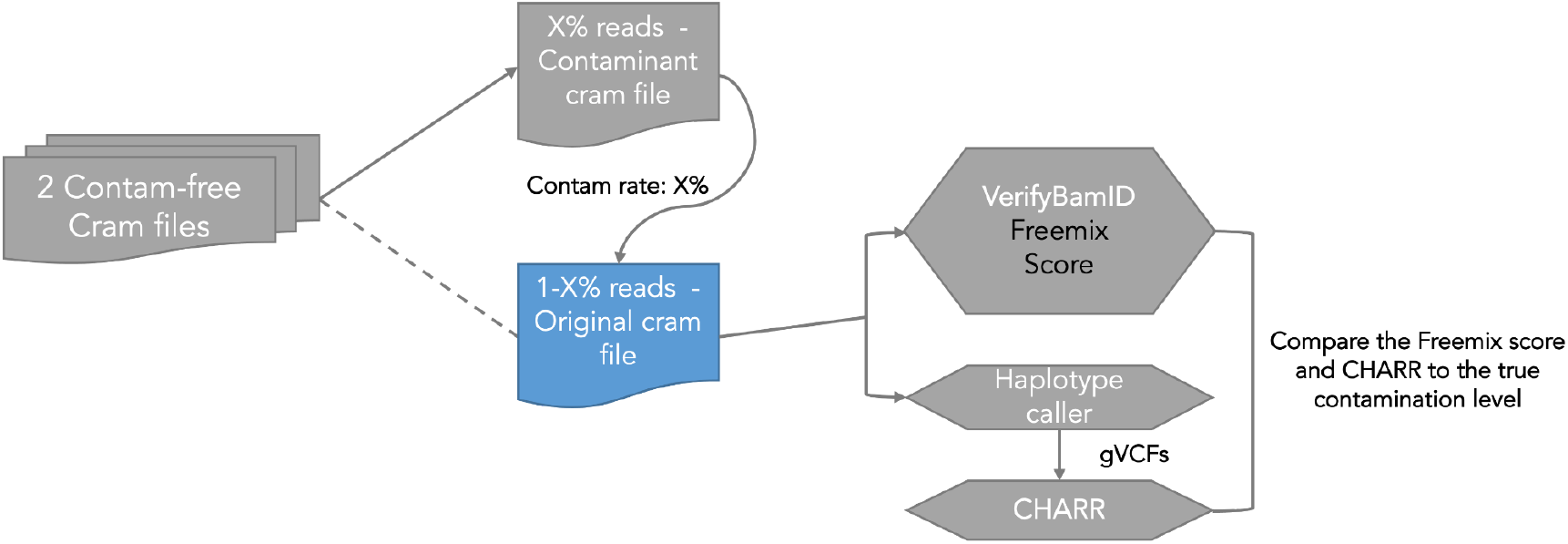
Pipeline for mixing two contamination-free samples with assigned contamination rate and implementing comparison of contamination rate produced by CHARR and Freemix Score.

To define a contamination rate of x%, we draw from a Bernoulli distribution with a probability x%, inserting a read from the contaminant on an observation of success, and inserting a read from the original sample otherwise. The newly-contaminated CRAM files are run through GATK HaplotypeCaller to generate gVCF files. This pipeline results in 30 samples x 5 contamination rates = 150 mixed samples, and we compute freemix scores on the 150 CRAM files and CHARR on the VDS joint called from the 150 merged gVCFs (Figure S14 to S15).

**Figure M4.**
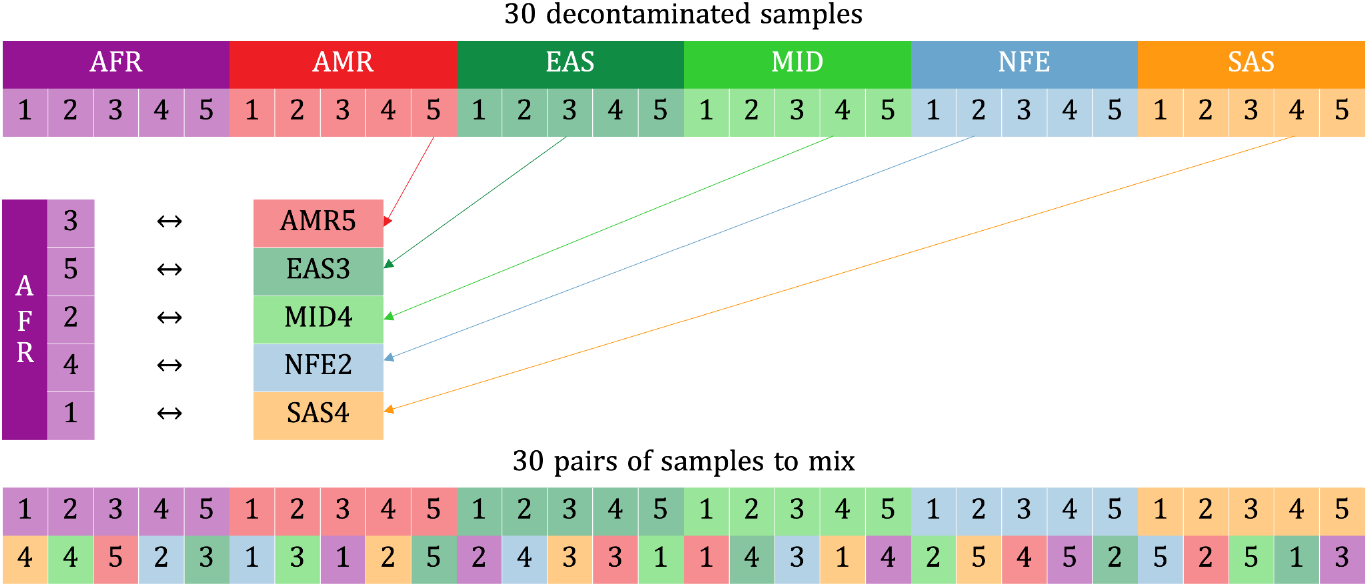
Strategy of randomly matching samples with different ancestry groups for the two-way simulation.

#### Mixing N contamination-free samples

As a single source of contamination may not be an appropriate scenario for large-scale sequencing projects, we further implemented an N-way mixing pipeline to produce simulated samples with contaminating reads from a pool of samples. Specifically, using the same 30 decontaminated samples and simulating under the same set of contamination rates, we contaminated the CRAM files of each sample with reads from any of the remaining 29 samples at the same five contamination rates, thus creating 150 simulated mixed files with more diverse sources of contamination (Figures S17 & S18).

For each read for each original sample, we decide whether to keep a read from the original sample by the same Bernoulli strategy as described in the two-way pipeline. If the random draw results in a contamination event, we randomly select a sample from the remaining 29 samples and insert the read from that selected sample. Once the simulated files were generated, we applied GATK (Ver 4.2.0.0) HaplotypeCaller to produce the gVCF files for each of the new mixed files and then implemented both CHARR and VerifyBamID to compute the contamination rates under these simulation scenarios.

## Acknowledgements

We thank Rasmus Nielsen for helpful discussions. This work was supported by the National Institutes of Health, including the National Human Genome Research Institute under award number U24HG011450 and National Institution of Mental Health under award number R37MH107649. The content is solely the responsibility of the authors and does not necessarily represent the official views of the National Institutes of Health.

## Resources

CHARR Analysis:

- https://github.com/atgu/CHARR

CHARR Implementations:https://github.com/HTGenomeAnalysisUnit/SCE-VCF

- https://github.com/hail-is/hail/blob/main/hail/python/hail/methods/qc.py#L1739

SVCR & VDS:

- https://hail.is/docs/0.2/vds/index.html#the-scalable-variant-call-representation-svcr

