## Supplemental Information for "CHARR efficiently estimates contamination from DNA sequencing data"

### Supplementary information for Lu et al., 2023


**Supplementary information for Lu et al., 2023** **1**

**Distribution of homozygous variants with reference reads** **3**

Figure S1 | The distribution of the number of high-quality homozygous variants with reference reads across samples in gnomAD v3. 3

Table S1 | The distribution of the number of high-quality homozygous variants with any reference reads (left) or at least 10% reference alleles (right) in gnomAD v3. 3

**Impact of variant selection on CHARR** **4**

Figure S2 | A comparison of the Freemix score and CHARR computed from autosomal, biallelic SNVs with GQ  20 and 20  DP  100 among gnomAD v3 WGS samples without and with a ref-AF cutoff (0-100% vs. 10-90%). The mean numbers of homozygous variants used for computing CHARR are 583,368 (left) and 463,668 (right). The black dashed line here represents y=x. 4

Figure S3 | Linear regression results from regressing CHARR computed from autosomal, biallelic SNVs with GQ  20 among gnomAD v3 WGS (panel A) and gnomAD v2 WES samples (panel B) samples on the Freemix score. The intercept (top), slope (middle), and r-squared (bottom) are shown for varying DP cutoffs (panel columns) and reference allele frequency cutoffs (x-axis). 6

Figure S4 | Linear regression results from regressing CHARR computed from autosomal, biallelic variants with GQ  20 and 20  DP  100 among gnomAD v3 WGS samples on the Freemix score varying reference allele frequency cutoffs with different variant sets. 7

**Sources of reference allele frequencies** **8**

Figure S5 | A comparison of Freemix score and CHARR computed from autosomal, biallelic SNVs with GQ  20 and 20  DP  100 among gnomAD v2 WES samples with reference allele frequencies from different sources. The mean numbers of homozygous variants used for computing CHARR are 805, 718, 575, 1012, 890, and 716, respectively for the six panels from left to right, top to bottom. The black dashed line here represents y=x. 9

Figure S6 | Linear regression results from regressing CHARR computed from autosomal, biallelic SNVs with GQ  20 and 20  DP  100 among gnomAD v2 WES samples on the Freemix score varying reference allele frequency cutoffs with reference allele frequencies from different sources. 9

**Mosaicism** **10**

Impact of known mosaicism genes on CHARR 10

Figure S7 | CHARR computed after removing three genes related to age (ASXL1, TET2, DNMT3A). The mean numbers of homozygous variants used for computing CHARR are 558573, 463641, 366549, 558604, 463668, and 366573, respectively for the six panels from left to right, top to bottom. The black dashed line here represents y=x. 11

Figure S8 | Linear regression results from regressing CHARR computed from autosomal, biallelic SNVs with GQ  20 and 20  DP  100 among gnomAD v3 WGS samples with three genes related to age (ASXL1, TET2, DNMT3A) removed on Freemix score. 11

Age vs. CHARR 12

Table S2 | The number of samples within each age group (for those with age defined). 12
Figure S9 | Comparison between age and CHARR computed from autosomal, biallelic SNVs with GQ  20, 20  DP 100 and 10% < reference AF < 90% for both gnomAD v2 WES samples (left) and v3 WGS samples (right). 12
**Versions of VerifyBamID** **13**
Figure S10 | Comparison of freemix scores computed from the old vs. new version of VerifyBamID. The black dashed line indicates y=x. 13
Figure S11 | Relationship between CHARR and Freemix scores computed from the old and new versions of VerifyBamID. The black dashed line here represents y=x. 14
**Simulation results** **15**
Figure S12 | 30 samples randomly selected from the HGDP database, including 5 samples from each of the 6 major genetic ancestry groups with their original contamination rates distributed approximately uniformly within each group. 15
Sample decontamination 15
Figure S13 | A comparison of the Freemix score and CHARR across the 30 selected samples from the HGDP dataset pre- and post-decontamination (circles and triangles, respectively). The cluster of decontaminated samples at the bottom of the figure demonstrates that the decontamination pipeline has effectively produced contamination-free short-read data. 16
Two-way sample mixing pipeline 16
Figure S14 | A comparison of Freemix score and CHARR for ~150 simulated 2-way mixed samples across five true contamination levels (by different colors). The black dashed lines represent y=x. CHARR from panel (A) is computed using allele frequencies from gnomAD, and that from panel (B) is computed using the corresponding allele frequencies of the single contaminant sample. 17
Figure S15 | A comparison of the ratio of the number of heterozygous and homozygous variants across the 150 2-way simulated samples. 18
N-way sample mixing pipeline 18
Figure S16 | A comparison of Freemix score and CHARR for ~150 simulated n-way mixed samples across five true contamination levels (indicated by different colors). The black dashed lines represent y=x, with grid lines at the true contamination rates. The columns indicate the sources of allele frequencies used for CHARR, and the rows indicate whether a 100% callrate filter has been applied. 20
Figure S17 | A comparison of Freemix score (green) and CHARR for ~150 simulated n-way mixed samples computed with different sources of allele frequencies (indicated by different colors). The black dashed lines represent the true contamination rate, which is also highlighted at the top of each panel. 20
Comparison between the N-way and 2-way results 21
Figure S18 | A comparison of Freemix score and CHARR for 300 simulated mixed samples across five true contamination levels, with 150 from the two-way mixing pipeline represented by the round dots and 150 from the n-way mixing pipeline represented by the triangular dots. CHARR is computed using the gnomAD reference allele frequencies. The black dashed lines represent the true contamination level. 21
**References** **22**

#### Distribution of homozygous variants with reference reads

We computed the number of homozygous alternate variants with reference reads above 0 across 59,765 gnomAD v3 WGS samples. The bimodality presented in the distribution of this metric is separated by the level of CHARR, which suggests samples with high contamination tend to have more homozygous variants that are infiltrated with reference reads (Figure S1).

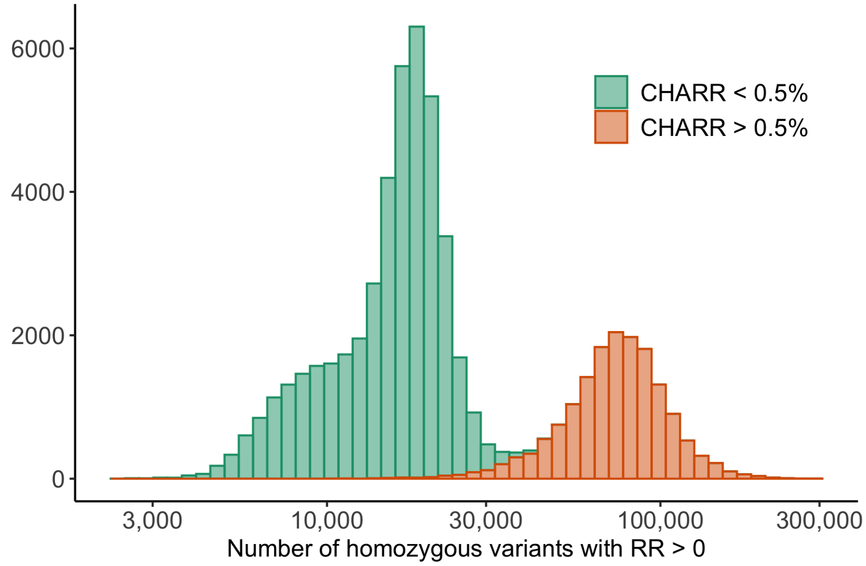


Figure S1 | The distribution of the number of high-quality homozygous variants with reference reads across samples in gnomAD v3.

Table S1 | The distribution of the number of high-quality homozygous variants with any reference reads (left) or at least 10% reference alleles (right) in gnomAD v3.

| **Homozygous variants with ≥ one reference read** | Number of samples | **Homozygous variants with at least 10% reference reads** | Number of samples |
|---|---|---|---|
| **> 0** | 59,765 | **> 0** | 59,765 |
| **> 10,000** | 54,849 | **> 30** | 48,460 |
| **> 20,000** | 26,446 | **> 50** | 28,950 |
| **> 30,000** | 16,376 | **> 100** | 3,564 |
| **> 100,000** | 2,665 | **> 500** | 169 |
| **> 200,000** | 40 | **> 1,000** | 102 |

#### Impact of variant selection on CHARR

We sought to assess how the choice of variants included impacts the CHARR score, particularly with respect to reference allele frequencies, sequencing depth, and variant type. For reference allele frequencies, we note that an extremely low reference allele frequency coupled with very low reference allele balance will return a very large value, inflating CHARR (Figure S2: left, ref_AF: (0%, 100%)). By filtering out variants with either reference allele or alternate allele frequency greater than 10%, the performance of CHARR substantially improves and highly aligns with the Freemix score, the contamination estimator from VerifyBamID (Figure S2: right, ref_AF: (10%, 90%)).

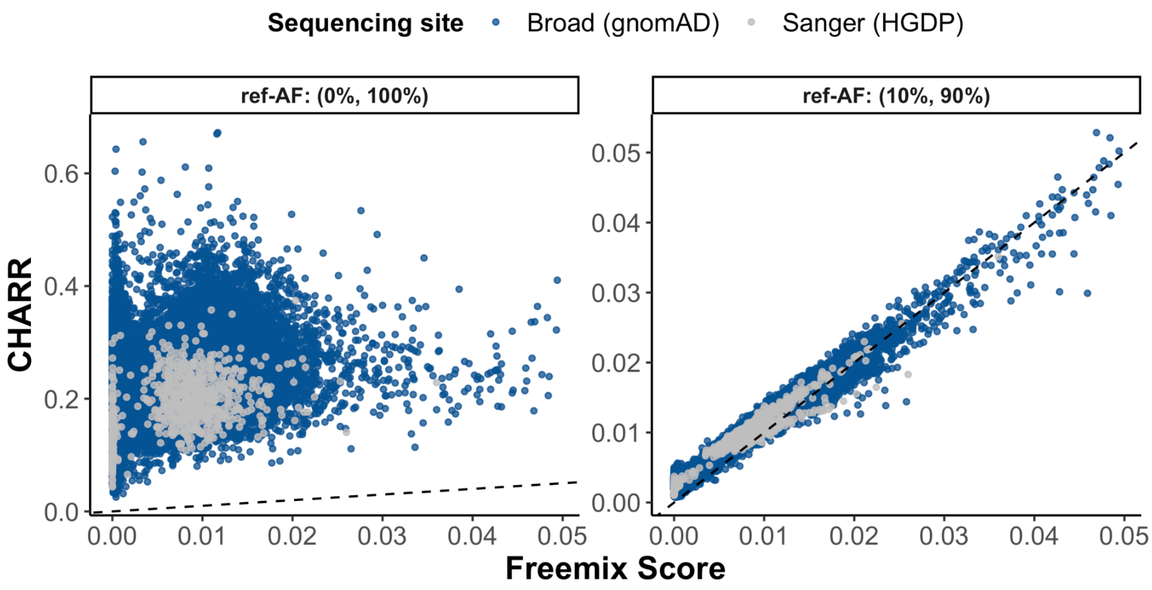


Figure S2 | A comparison of the Freemix score and CHARR computed from autosomal, biallelic SNVs with GQ ≥ 20 and 20 ≤ DP ≤ 100 among gnomAD v3 WGS samples without and with a ref-AF cutoff (0-100% vs. 10-90%). The mean numbers of homozygous variants used for computing CHARR are 583,368 (left) and 463,668 (right). The black dashed line here represents y=x.

We systematically investigated the impact of sequencing depth and allele frequency on the performance of CHARR. We performed a linear regression on CHARR against the Freemix scores with three different cutoff schemes of sequencing depth (DP) applied: 1) No DP cutoff; 2)

10 $\leq$ DP $\leq$ 100; 3) 20 $\leq$ DP $\leq$ 100 and across four reference allele frequency (ref-AF) cutoffs: 1) 1% $\leq$ ref-AF $\leq$ 99%; 2) 5% $\leq$ ref-AF $\leq$ 95%; 3) 10% $\leq$ ref-AF $\leq$ 90%; 4) 20% $\leq$ ref-AF $\leq$ 80%. We evaluate the performance of CHARR based on three criteria: 1) the magnitude of the intercept; 2) how much the slope deviates from 1; 3) the value of the R-squared statistics.

Among all the filtering schemes on sequencing depth, we achieve the best performance with 20 $\leq$ DP $\leq$ 100, which has the highest R-squared, slopes closer to 1, and a relatively small intercept among the gnomAD v3 WGS samples (Figure S3A). Similarly, for the results from the 102,063 release WES samples from gnomAD v2, though the performance of CHARR declines with fewer homozygous variants, it still maintains a reasonably high correlation with the Freemix score for 20 $\leq$ DP $\leq$ 100 (Figure S3B). This comparison demonstrates that it is also feasible to downsample the WGS data to a subset of variants, e.g., the exonic regions, while computing CHARR, which can achieve reasonable accuracy at a reduced computational cost.

As we impose a stricter cutoff on reference allele frequency, the intercept gets closer to zero, and the value of R-squared increases to over 0.96 while the slope slightly decreases. In practice, we can achieve a high correlation with any cutoff at or above 5%, and a similar trend is observed for both gnomAD v3 WGS and gnomAD v2 WES samples. Therefore, we recommend applying a symmetric cutoff on the reference allele frequency of at least 5% to maintain reasonable accuracy of contamination level. These results suggest that our method is robust to choice of depth and allele frequency thresholds, as we can obtain satisfactory results at any allele frequency cutoff above 5% (Figure S3).

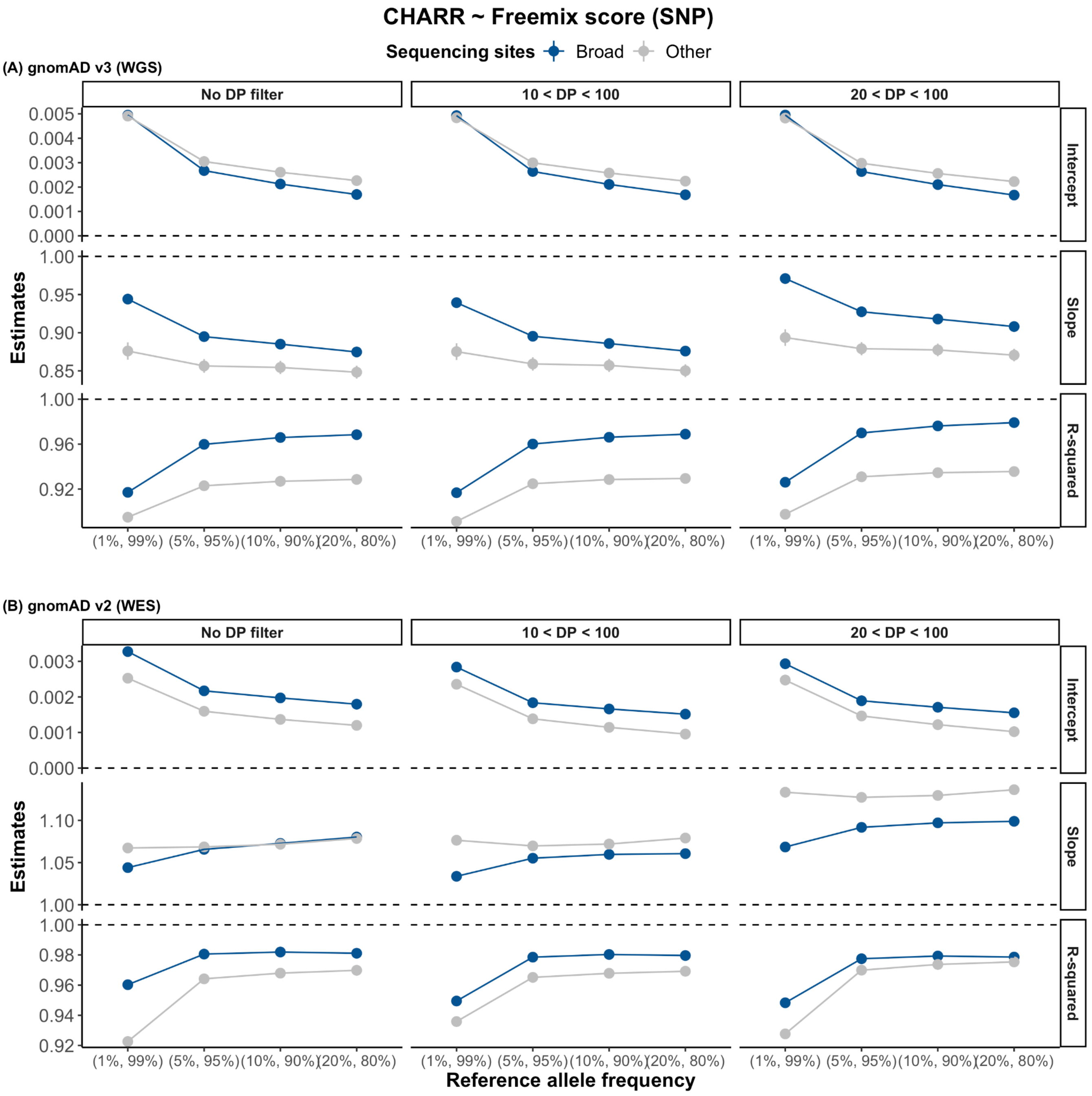


Figure S3 | Linear regression results from regressing CHARR computed from autosomal, biallelic SNVs with GQ ≥ 20 among gnomAD v3 WGS (panel A) and gnomAD v2 WES samples (panel B) samples on the Freemix score. The intercept (top), slope (middle), and r-squared (bottom) are shown for varying DP cutoffs (panel columns) and reference allele frequency cutoffs (x-axis).

We also test CHARR with various variant categories: 1) SNPs + INDELs, 2) INDELs, and 3) SNPs within the gnomAD v3 WGS samples. Under our preferred threshold combination of 20 ≤ DP ≤ 100 and 10% ≤ ref-AF ≤ 90%, all three scenarios demonstrate good alignments between CHARR and the Freemix score (R-squared > 90% for most of the cases). CHARR estimated from the INDELs shows a modest deflation of the slope compared to SNPs or the combined score, but maintains a high R-squared of over 0.95 (Figure S4). Therefore, we show further evidence here of the robustness of CHARR that it can also be computed from INDELs, a much smaller subset of the variants from the WGS data.

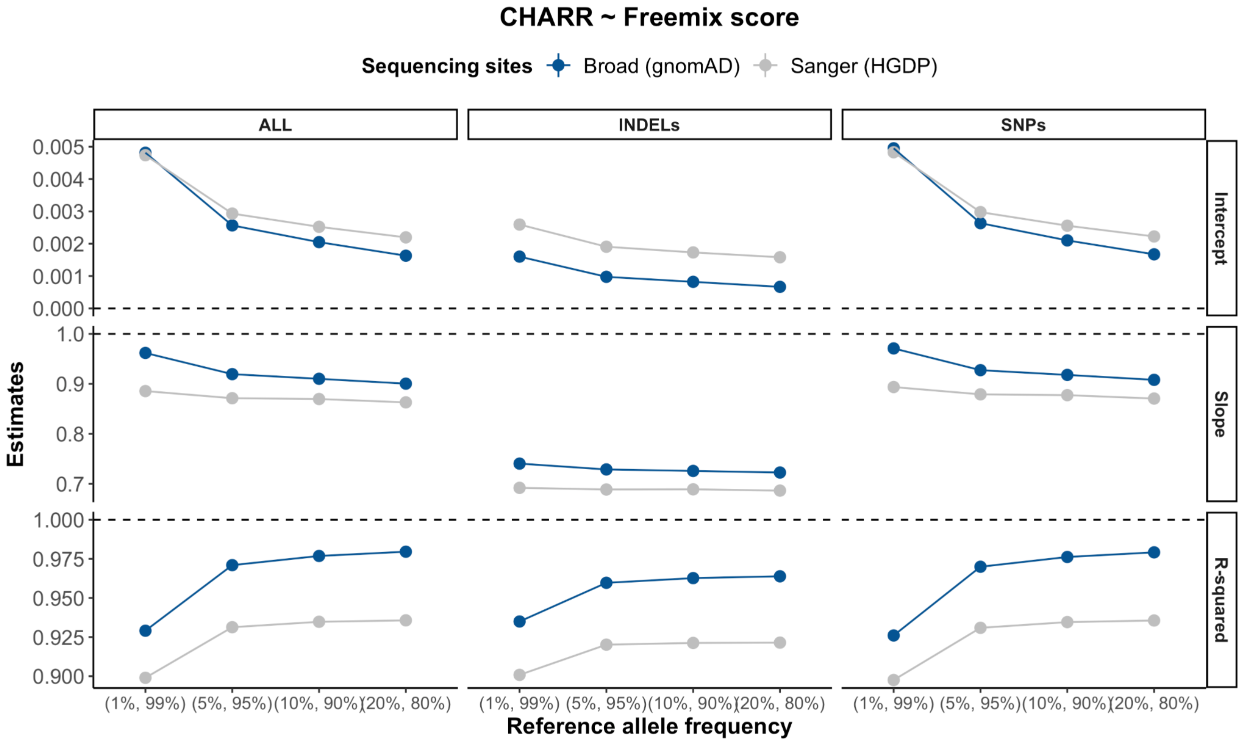


Figure S4 | Linear regression results from regressing CHARR computed from autosomal, biallelic variants with GQ ≥ 20 and 20 ≤ DP ≤ 100 among gnomAD v3 WGS samples on the Freemix score varying reference allele frequency cutoffs with different variant sets.

#### Sources of reference allele frequencies

An important element to computing CHARR involves the accurate estimation of reference allele frequencies (ref-AF) that best reflect the source of contaminating samples, which is inherently unknown. In particular, this could be obtained within the callset as an approximation of the diversity of contaminating samples from the same sequencing environment; however, this is likely to overestimate allele frequencies of rare variants. Alternatively, these metrics could be estimated from external sources, which may be a better approach when the target callset is small. To investigate the impact of applying different sources of global ref-AF, i.e., ref-AF from WGS and WES, we computed CHARR on the gnomAD v2 WES data with both sources of ref-AFs and compared their performances. The correlations between CHARR and the Freemix score are very similar in these two scenarios (Figure S5). Using ref-AF computed from the WES data slightly improves the results from the linear regression (Figure S6), but using the WGS ref-AF also results in a CHARR with reasonably high quality, where the linear regression results show r-squared > 0.96 and slope closer to 1 than using the WES ref-AF when ref-AF cutoffs are 5% or above. Therefore, it is feasible to use a reference allele frequency from a large reference dataset, which is also demonstrated in the simulation results that using gnomAD AF produces CHARR with equivalently good accuracy as VerifyBamID (see below; Figure S17).

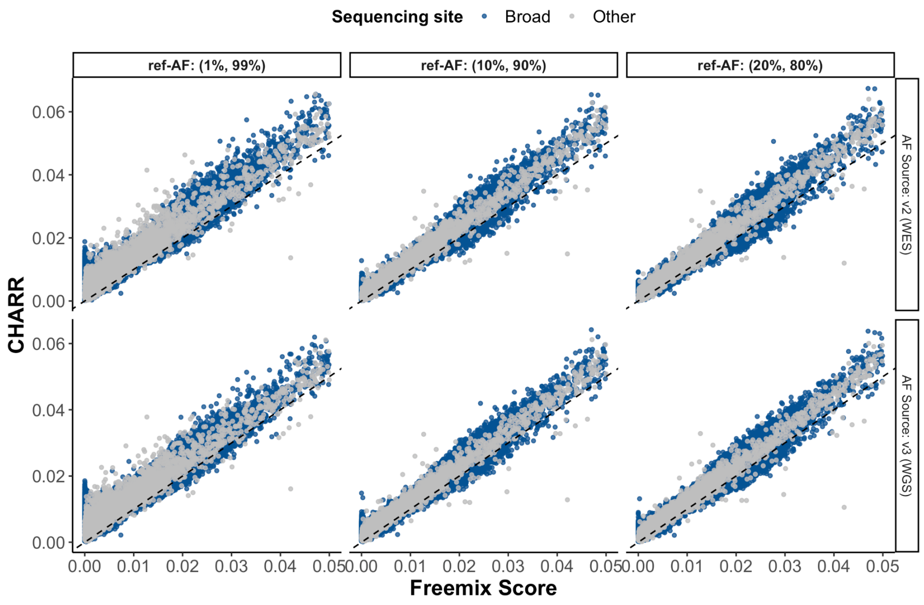


Figure S5 | A comparison of Freemix score and CHARR computed from autosomal, biallelic SNVs with GQ ≥ 20 and 20 ≤ DP ≤ 100 among gnomAD v2 WES samples with reference allele frequencies from different sources. The mean numbers of homozygous variants used for computing CHARR are 805, 718, 575, 1012, 890, and 716, respectively for the six panels from left to right, top to bottom. The black dashed line here represents y=x.

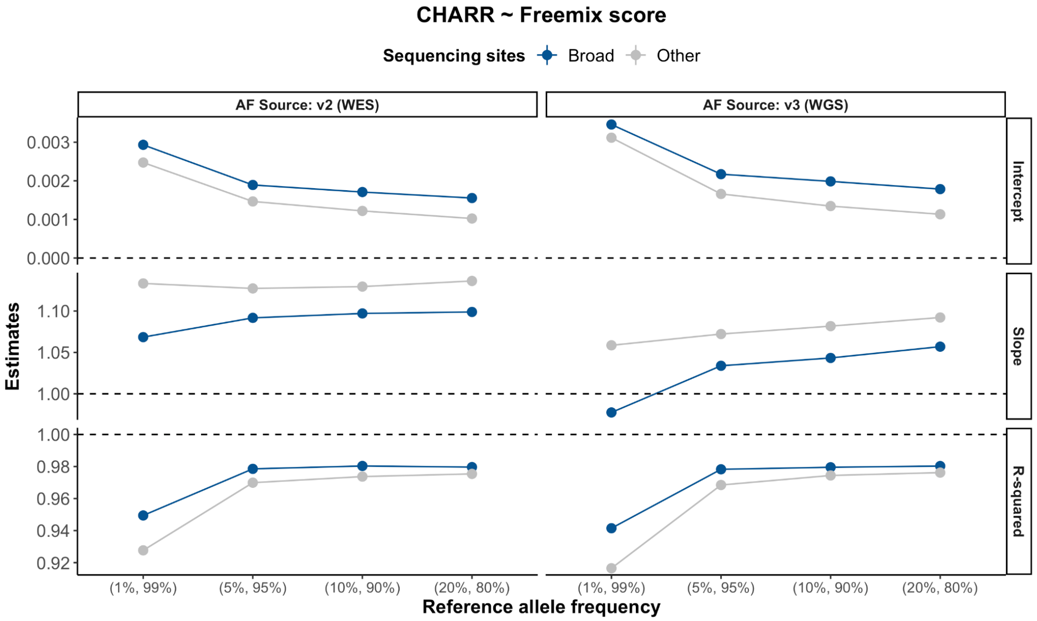


Figure S6 | Linear regression results from regressing CHARR computed from autosomal, biallelic SNVs with GQ ≥ 20 and 20 ≤ DP ≤ 100 among gnomAD v2 WES samples on the Freemix score varying reference allele frequency cutoffs with reference allele frequencies from different sources.

#### Mosaicism

Genetic mosaicism is defined as the presence of two or more genetically different sets of cells in a single individual, which can act as an additional source of the imbalanced reference allele depth. Previous studies have demonstrated that the occurrence of mosaicism is positively related to age (Jacobs et al., 2012). Therefore, to investigate this potential factor, we excluded three genes where mutations were previously identified to be associated with age (Xie et al., 2014) and recomputed CHARR with the remaining variants. There is no noticeable difference in the performance of CHARR before and after the removal of the three mosaicism-related genes (Figures S7 & S8). Meanwhile, we examine the relationship between CHARR and age, which positively correlates with the occurrence of mosaicism. We find no evidence of correlation across all age groups within both the WGS and the WES data (Figure S9).

##### *Impact of known mosaicism genes on CHARR*

Though DNA sample contamination accounts for many of the unexpected observations of reference reads within the homozygous alternate variants, another possible explanation of such infiltration could be mosaicism within the target individual. To determine this, we compared CHARR before and after removing three genes that were previously found to be associated with mosaicism with age, *ASXL1*, *TET2*, and *DNMT3A*. The results demonstrate that these three genes do not have a compelling impact on the overall estimation of contamination level (Figure S7). With all SNPs included in the computation, CHARR reaches a reasonable estimate of both the slopes and intercepts from the linear regression while maintaining a similar level of the R-squared statistics (Figure S8).

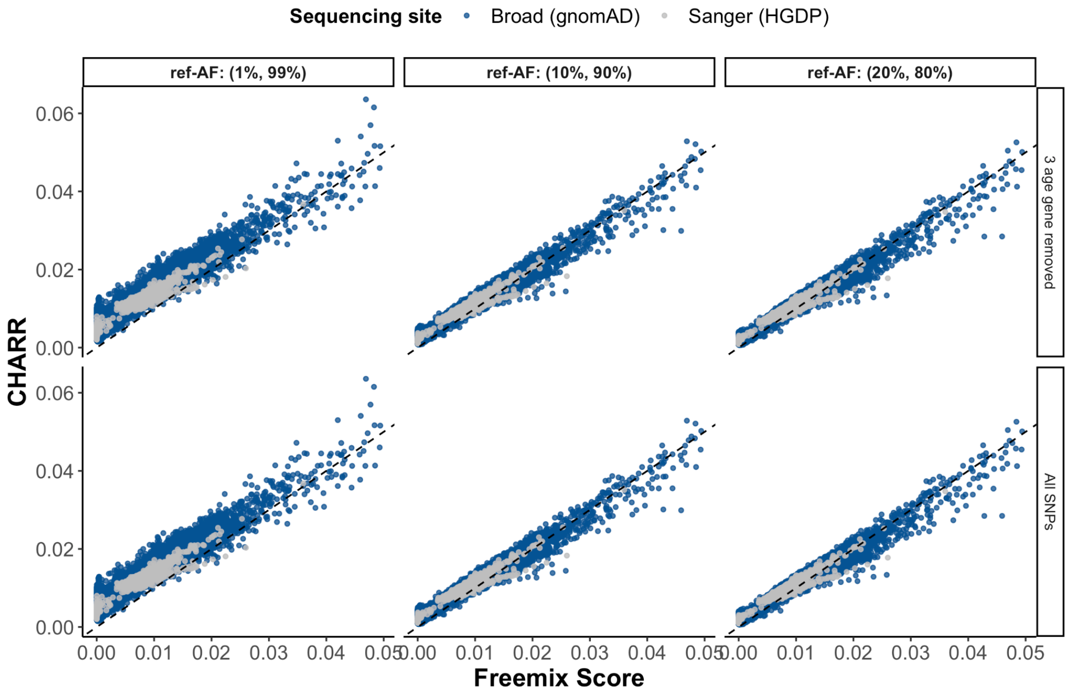


Figure S7 | CHARR computed after removing three genes related to age (*ASXL1, TET2, DNMT3A*). The mean numbers of homozygous variants used for computing CHARR are 558573, 463641, 366549, 558604, 463668, and 366573, respectively for the six panels from left to right, top to bottom. The black dashed line here represents y=x.

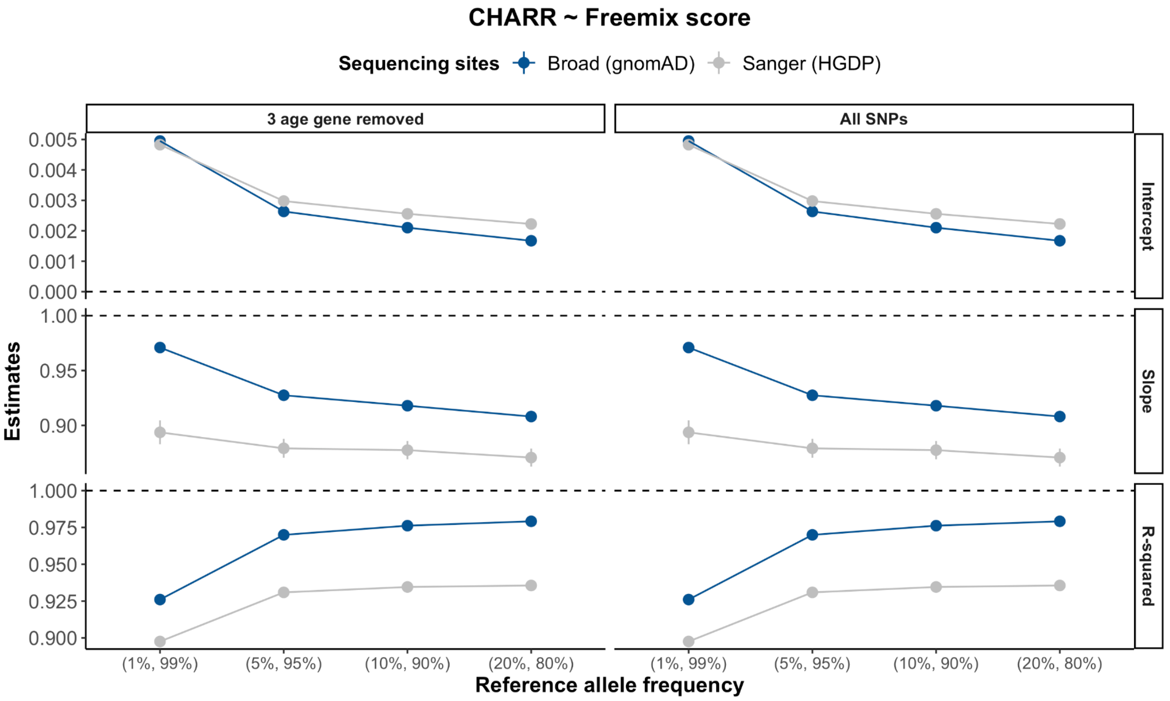


Figure S8 | Linear regression results from regressing CHARR computed from autosomal, biallelic SNVs with GQ ≥ 20 and 20 ≤ DP ≤ 100 among gnomAD v3 WGS samples with three genes related to age (*ASXL1, TET2, DNMT3A*) removed on Freemix score.

*Age vs. CHARR*

To further understand the potential influence of mosaicism, we examine the relationship between age (Table S2) and CHARR computed from autosomal, biallelic SNVs with $20 \leq DP \leq 100$ and 10% < reference AF < 90% in gnomAD v3 and v2. We find no correlation between the two metrics, suggesting no apparent impact on CHARR from mosaicism (Figure S9).

Table S2 | The number of samples within each age group (for those with age defined).

| **Age Bin** | **0-10** | **10-20** | **20-30** | **30-40** | **40-50** | **50-60** | **60-70** | **70-80** | **> 80** |
|---|---|---|---|---|---|---|---|---|---|
| gnomAD v3 | 158 | 1,407 | 2,422 | 2,634 | 4,910 | 7,114 | 5,331 | 2,506 | 392 |
| gnomAD v2 | 129 | 787 | 788 | 2,555 | 7,224 | 10,195 | 7,503 | 4,221 | 1,451 |

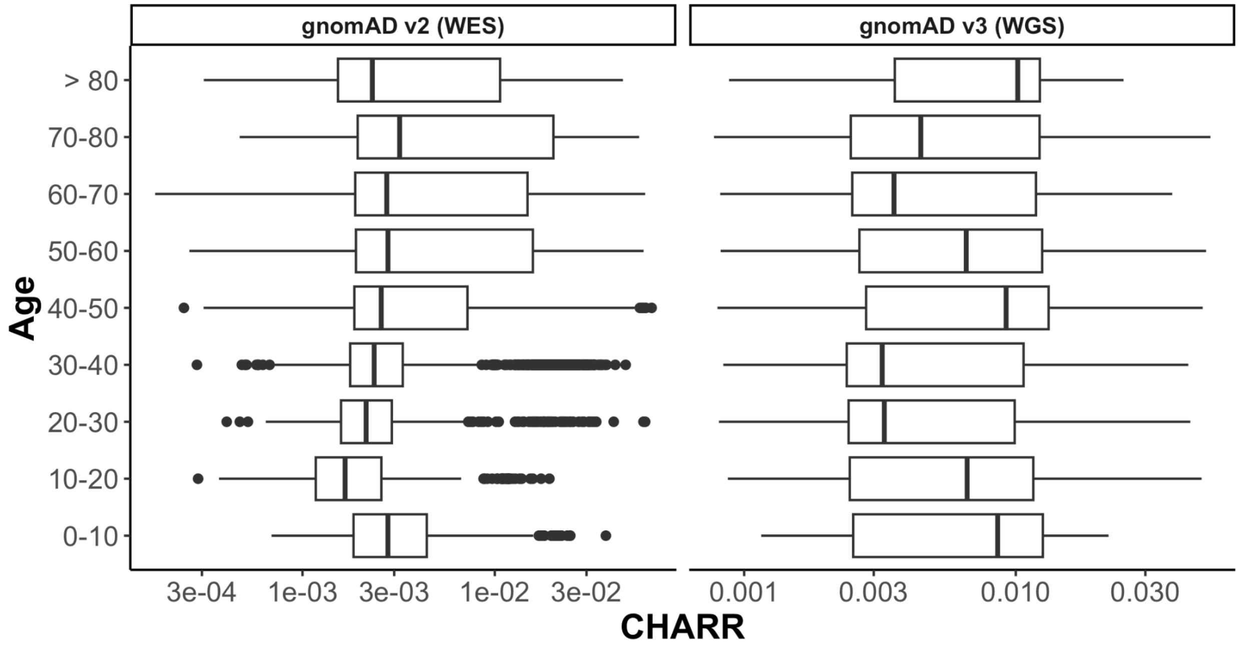


Figure S9 | Comparison between age and CHARR computed from autosomal, biallelic SNVs with $GQ \geq 20$, $20 \leq DP \leq 100$ and 10% < reference AF < 90% for both gnomAD v2 WES samples (left) and v3 WGS samples (right).

#### Versions of VerifyBamID

We also identified differences in estimates produced by different configurations of the VerifyBamID task used to calculate rates of cross-sample contamination. In the course of processing gnomAD data, the pipeline to run VerifyBamID was updated. The earlier version of the task ran VerifyBamID 1.0.1, which had been previously found to produce inflated contamination estimates in some admixed samples. The resource files were then updated to the 1000 Genomes 100,000 sites version from 1.0.6. The contamination rates computed from the older VerifyBamID are slightly deflated compared to newer versions (Figure S10). Across all 948 HGDP samples, we recomputed using the latest VerifyBamID and compared the Freemix score from the early version and the latest version of VerifyBamID to our CHARR estimator. We find that while the previous version is somewhat inflated, the latest version is more concordant with our CHARR estimator with a slope estimate close to 1 (Figure S11).

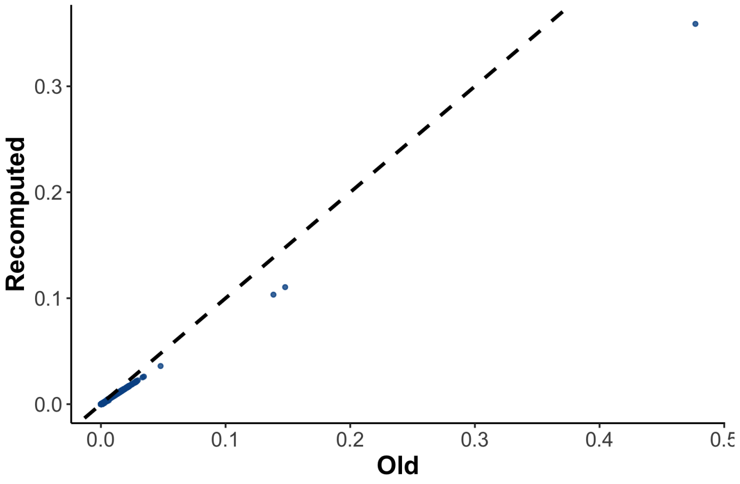


Figure S10 | Comparison of freemix scores computed from the old vs. new version of VerifyBamID. The black dashed line indicates y=x.

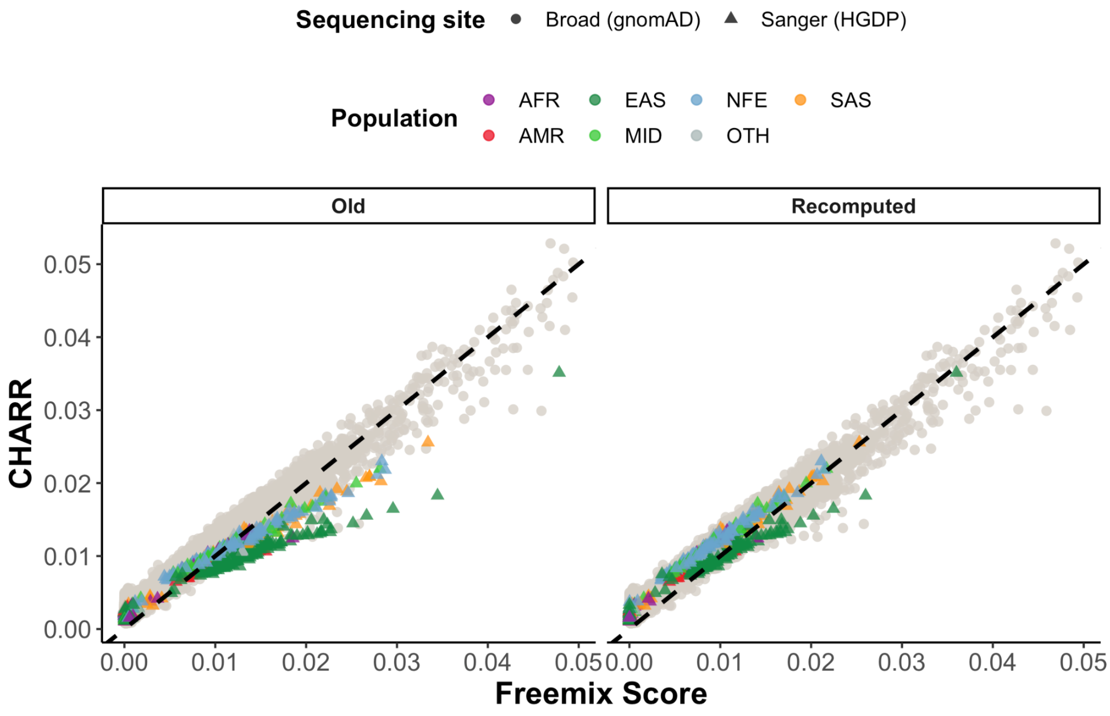


Figure S11 | Relationship between CHARR and Freemix scores computed from the old and new versions of VerifyBamID. The black dashed line here represents y=x.

#### Simulation results

We randomly selected 30 samples from the HGDP dataset evenly distributed across each genetic ancestry group and the spectrum of contamination levels (Figure S12). As described in Methods, we generated “decontaminated” samples, and produced simulated samples by mixing the post-decontamination CRAM files of these samples under two potential contamination scenarios, and recomputed contamination using VerifyBamID and CHARR.

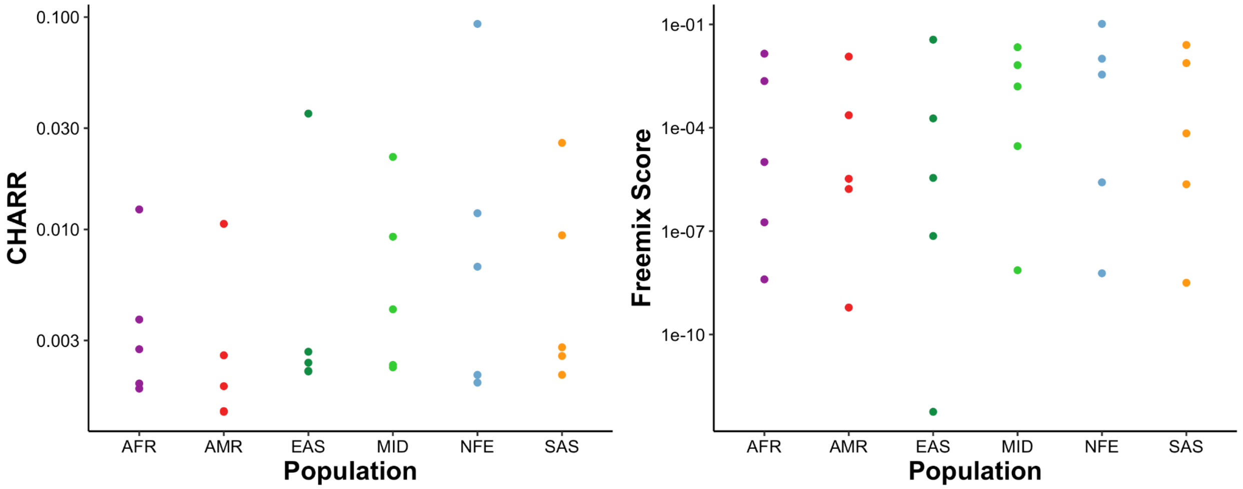


Figure S12 | 30 samples randomly selected from the HGDP database, including 5 samples from each of the 6 major genetic ancestry groups with their original contamination rates distributed approximately uniformly within each group.

##### *Sample decontamination*

The decontamination pipeline effectively decreases both the CHARR and Freemix scores (Figure S13). Notably, some samples which had extremely low Freemix scores prior to decontamination (< $10^{-9}$) resulted in slightly increased scores (~ $10^{-9}$); however, all samples clustered around a minimal value for both metrics (freemix $10^{-9}$ to $10^{-8}$, and CHARR ~$10^{-4}$), indicating that the pipeline has successfully created a set of samples free from any detectable contamination.

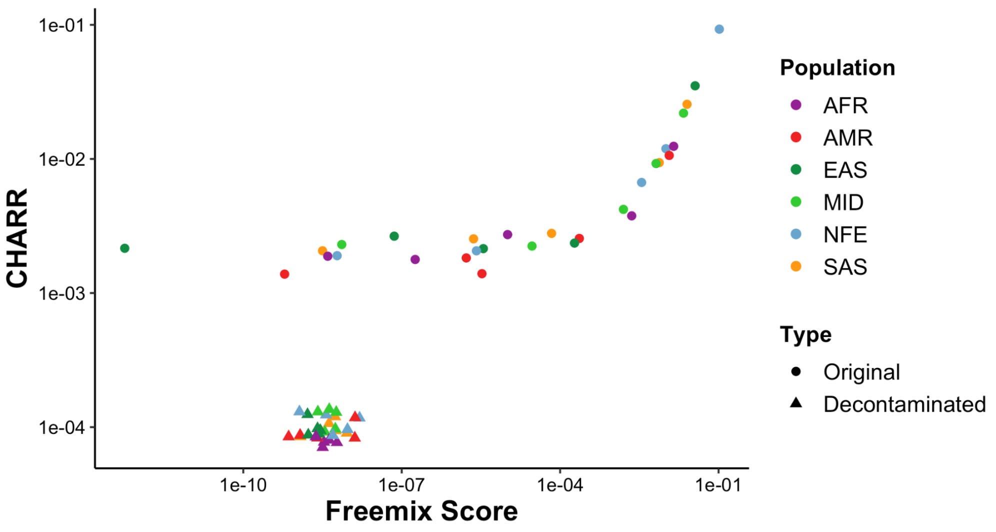


Figure S13 | A comparison of the Freemix score and CHARR across the 30 selected samples from the HGDP dataset pre- and post-decontamination (circles and triangles, respectively). The cluster of decontaminated samples at the bottom of the figure demonstrates that the decontamination pipeline has effectively produced contamination-free short-read data.

*Two-way sample mixing pipeline*

We mix 30 random pairs of samples to manually create simulated samples with contamination rates of 0.5%, 1%, 2%, 5%, and 10% (see Methods) and compute CHARR and Freemix scores for the 30 samples x 5 contamination rates = 150 mixed samples. At low contamination levels (0.5% and 1%), CHARR more accurately reflects the true contamination rate. However, as the contamination level further increases to 5-10%, though still increasing linearly, CHARR underestimates the true contamination rate. Using an allele frequency computed from the actual contaminating sample (“local”) produces slightly better results but still deviates from the truth values (Figure S14).

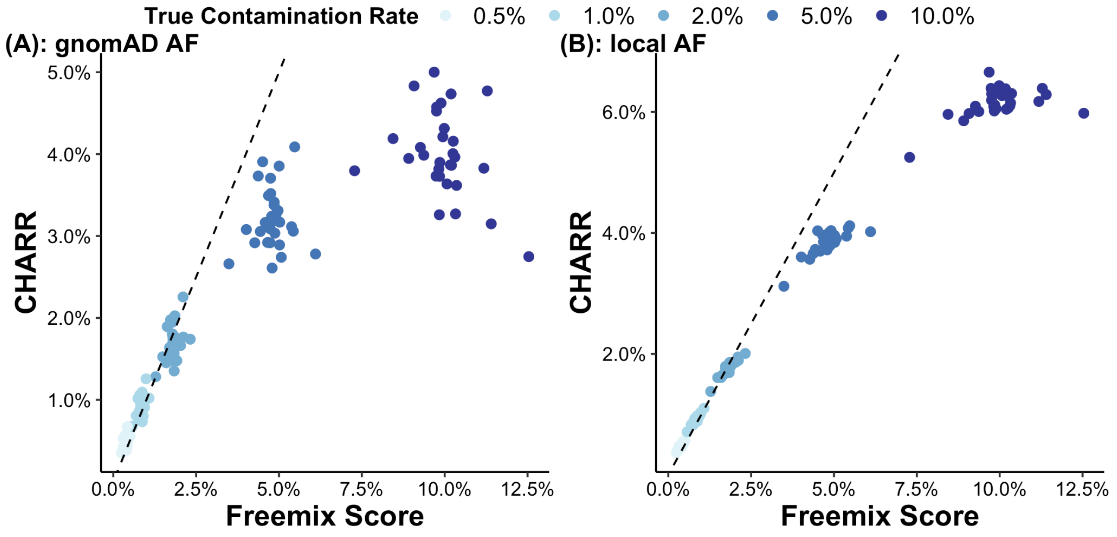


Figure S14 | A comparison of Freemix score and CHARR for ~150 simulated 2-way mixed samples across five true contamination levels (by different colors). The black dashed lines represent y=x. CHARR from panel (A) is computed using allele frequencies from gnomAD, and that from panel (B) is computed using the corresponding allele frequencies of the single contaminant sample.

We hypothesized that the deflation in CHARR may be due to the simulation strategy introducing an unlikely scenario of contamination, where it may be unreasonable to observe a single sample contaminated by a single other sample at rates of 5-10%. Notably, at 30x coverage and 5% contamination from a single sample, at a site that is homozygous alternate in the target sample and heterozygous in the contaminant, the expected reference allele balance in the simulated samples spans across 0% to 20%. For most variant calling algorithms (including GATK's HaplotypeCaller), the presence of ~7-10% reference alleles at a site will convert the site to heterozygous, decreasing the number of homozygous variants that would contribute substantially to CHARR, resulting in a deflation. Indeed, we observe a higher ratio of heterozygous to homozygous variants for samples simulated at a contamination rate of 10% and a slight inflation of the ratios in those with a contamination rate of 5% (Figure S15). Thus, we recommend computing this ratio as a part of sample quality control to identify samples that are highly contaminated by this mechanism.

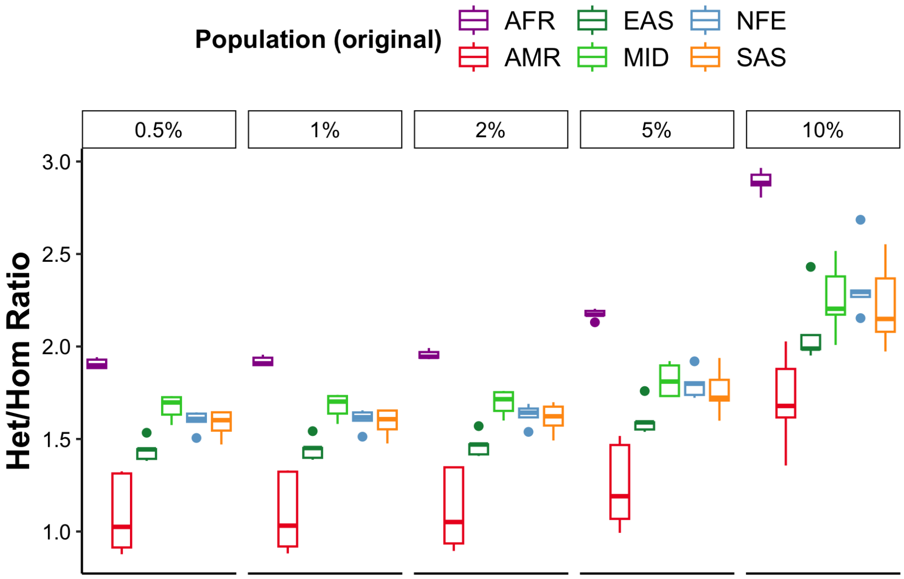


Figure S15 | A comparison of the ratio of the number of heterozygous and homozygous variants across the 150 2-way simulated samples.

*N-way sample mixing pipeline*

We next produced simulated samples with contaminants from a random combination of the 30 decontaminated samples under true contamination rates of 0.5%, 1%, 2%, 5%, and 10% (see Methods) and compared CHARR and Freemix scores of the 30 samples x 5 rates = 150 samples under multiple scenarios. We computed CHARR with different sources of reference allele frequencies, which are generated from the 76,156 gnomAD v3 release samples and the 30 decontaminated samples as described in the previous section. Within each cohort, we further filtered variants to those with callrate = 100% to narrow down our selection of variants with higher confidence, better approximating the true allele frequency.

Overall, CHARR is highly consistent with the Freemix score ($r > 0.997$, $p < 1\times10^{-100}$) regardless of the source of reference allele frequencies being used (Figure S16). For CHARR computed from gnomAD AF, we see a high concordance with VerifyBamID (Figure S16); however, both metrics are slightly deflated compared to the true simulated contamination rate.

Using the exact allele frequency from the known contaminating samples ("local") with a 100% callrate filter, which is the exact frequency of the contaminant cohort, CHARR performs most consistently and accurately among all estimators and across most of the true contamination rates (Figure S16-S17). Despite the generally high accuracy, at the 10% contamination rate, CHARR using the best possible frequency (callrate filtered local AF) still underestimates contamination by about 10% of the true value, which could be due to a continued deflation due to genotype conversion to heterozygotes (see above).

This result indicates that the selection of reference allele frequencies is a critical factor in computing CHARR, and highlights a limitation of our simulation study, that the simulated sample size is relatively small (30 decontaminated samples) compared to large sample sizes in real-world cohorts (hundreds of thousands of samples), which might introduce bias and suggest a future broader simulation approach. In practice, this also suggests that the best approach would be to use the precise allele frequencies of the contaminating samples; however, these are inherently unknown. However, using the original gnomAD v3 allele frequency returns comparable results to VerifyBamID, with a slight deflations at higher contamination rates (>5%). Therefore, using allele frequencies from gnomAD becomes a feasible alternative approach in that it is convenient and quick to obtain as shown in both the real (Figure 2) and simulated results (Figure S17).

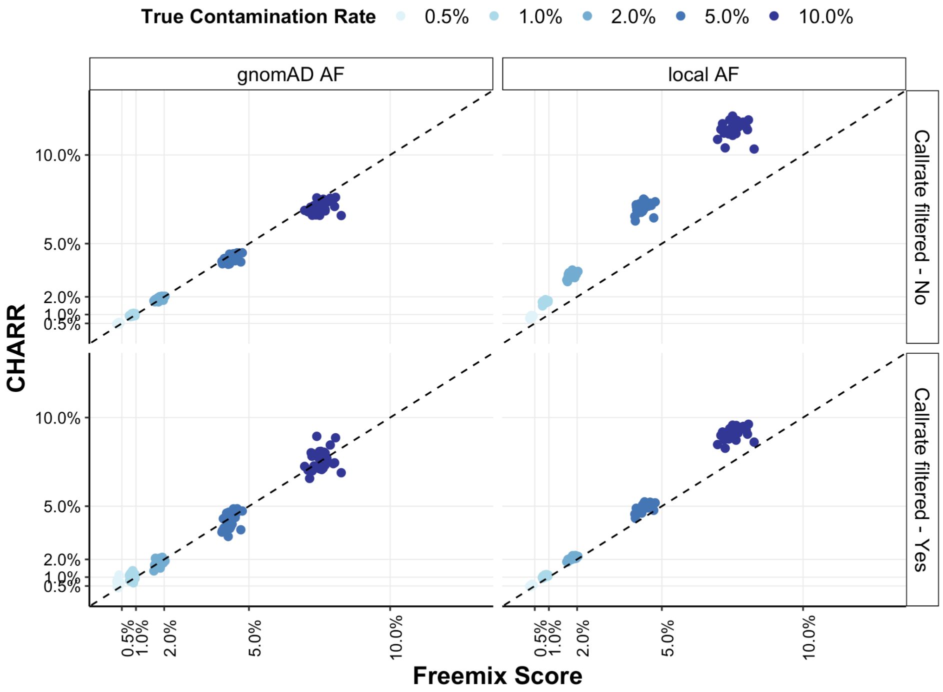


Figure S16 | A comparison of Freemix score and CHARR for ~150 simulated n-way mixed samples across five true contamination levels (indicated by different colors). The black dashed lines represent y=x, with grid lines at the true contamination rates. The columns indicate the sources of allele frequencies used for CHARR, and the rows indicate whether a 100% callrate filter has been applied.

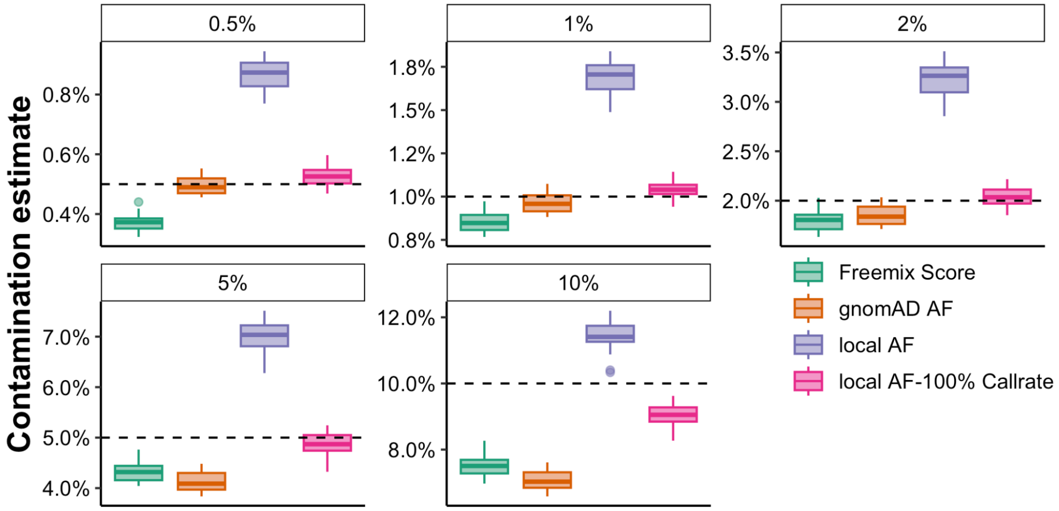


Figure S17 | A comparison of Freemix score (green) and CHARR for ~150 simulated n-way mixed samples computed with different sources of allele frequencies (indicated by different colors). The black dashed lines represent the true contamination rate, which is also highlighted at the top of each panel.

*Comparison between the N-way and 2-way results*

The results of CHARR and freemix are highly consistent between the 2-way and the n-way results when the true contamination rate is below 5%, with a slightly higher variance for both metrics in the 2-way simulation regime (Figure S18). When the contamination rate is above 5%, CHARR is deflated in the 2-way regime but consistent with Freemix for the n-way regime.

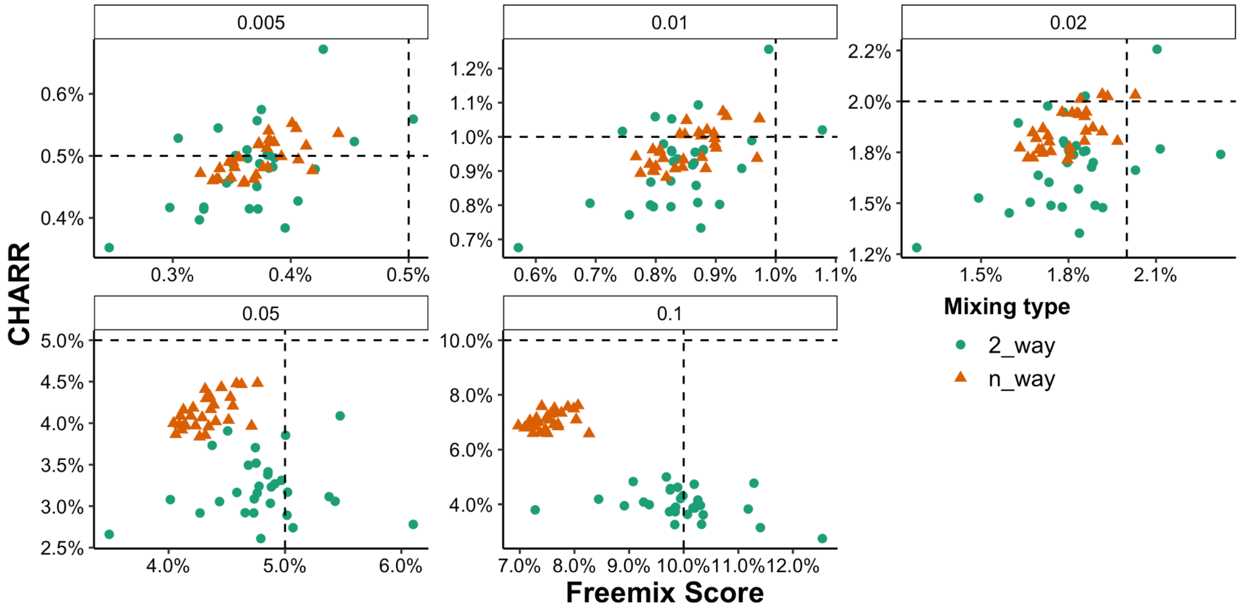


Figure S18 | A comparison of Freemix score and CHARR for 300 simulated mixed samples across five true contamination levels, with 150 from the two-way mixing pipeline represented by the round dots and 150 from the n-way mixing pipeline represented by the triangular dots. CHARR is computed using the gnomAD reference allele frequencies. The black dashed lines represent the true contamination level.

It is unclear which of these two is a more realistic scenario in real-world data. However, we note the existence of high CHARR (>5%) samples in gnomAD data (Figure 2) and their consistency with Freemix at these high contamination levels. If the 2-way mixing scenario were common, we would observe a deflation of CHARR compared to freemix at these levels in real data. However, as we see here that CHARR is more consistent with Freemix in the n-way regime, this suggests that the n-way regime is likely the more common scenario for real data.
